# Structural Insights into the Roles of PARP4 and NAD^+^ in the Human Vault Cage

**DOI:** 10.1101/2024.06.27.601040

**Authors:** Jane E. Lodwick, Rong Shen, Satchal Erramilli, Yuan Xie, Karolina Roganowicz, Anthony A. Kossiakoff, Minglei Zhao

## Abstract

Vault is a massive ribonucleoprotein complex found across Eukaryota. The major vault protein (MVP) oligomerizes into an ovular cage, which contains several minor vault components (MVCs) and is thought to transport transiently bound “cargo” molecules. Vertebrate vaults house a poly (ADP-ribose) polymerase (known as PARP4 in humans), which is the only MVC with known enzymatic activity. Despite being discovered decades ago, the molecular basis for PARP4’s interaction with MVP remains unclear. In this study, we determined the structure of the human vault cage in complex with PARP4 and its enzymatic substrate NAD^+^. The structures reveal atomic-level details of the protein-binding interface, as well as unexpected NAD^+^-binding pockets within the interior of the vault cage. In addition, proteomics data show that human vaults purified from wild-type and PARP4-depleted cells interact with distinct subsets of proteins. Our results thereby support a model in which PARP4’s specific incorporation into the vault cage helps to regulate vault’s selection of cargo and its subcellular localization. Further, PARP4’s proximity to MVP’s NAD^+^-binding sites could support its enzymatic function within the vault.

## Introduction

The vault particle was identified more than three decades ago and has been found within most eukaryotic clades [1], yet its molecular function is poorly understood. The particle’s primary structural element is known as the vault cage, an elliptical protein assembly measuring about 70 by 40 nm in dimension. In every species analyzed so far, the cage contains 78 copies of the major vault protein (MVP), which coalesce into two symmetrical, domed, 39mer halves. Three constitutive minor vault components (MVCs) have been identified in the cage in vertebrates: poly (ADP-ribose) polymerase 4 (PARP4), telomerase component 1 (TEP1), and a class of small non-coding RNAs known as vault RNAs (vtRNAs). Vault activity has been implicated in the innate immune response [2, 3], tumor progression & chemotherapeutic drug resistance [4, 5], and metabolic regulation [6, 7]. The vault cage structure has inspired hypotheses that vault acts as a signaling scaffold, reaction crucible, or transport module for endogenous, transiently internalized molecules known as its “cargo.” Indeed, MVP has been implicated in the signaling activity and relocalization of specific macromolecules in response to different cellular perturbations [6], [8–10]. However, its mechanism at the molecular level, including how it selects cargo molecules and their destinations, has not been elucidated. We hypothesized that some of these important regulatory roles might be performed by the MVCs, whose functions are not yet clear. In this study, we focus on PARP4, which is the only MVC with known catalytic activity.

PARP4 is a member of the PARP superfamily of enzymes, which consume the critical metabolite NAD^+^ to deposit ADP-ribose (ADPr) modifications onto macromolecular substrates. The biological function of PARP proteins is driven in large part by their “targeting” domains, which determine their subcellular localization and, consequently, the molecules they modify [11]. PARP4’s specific recruitment to the interior of the vault cage via its <u>M</u>VP-<u>In</u>teraction or “MINT” domain (Fig. 1a) implies that it could regulate vault activity by modifying MVP, other minor components, or encapsulated vault cargo. To this end, PARP4 contains an unusual combination of domains involved in macromolecular interactions, including VWA, VIT, and BRCT (Fig. 1a), which could facilitate direct interactions between PARP4 and vault’s protein or nucleic acid cargo. In addition, there is broad interest in engineering vault to deliver biomolecules into cells. Fusing these molecules to the MINT domain has proved to be an effective way to package them into the vault cage [12], generating interest in mutating the MINT domain to orchestrate optimal molecular loading and release [13]. These properties motivated us to determine the structural basis of PARP4’s recruitment to the vault cage, the impact of adding PARP4’s substrate NAD^+^ to the complex, and the effect of PARP4’s presence on the vault interactome.

**Fig 1.**
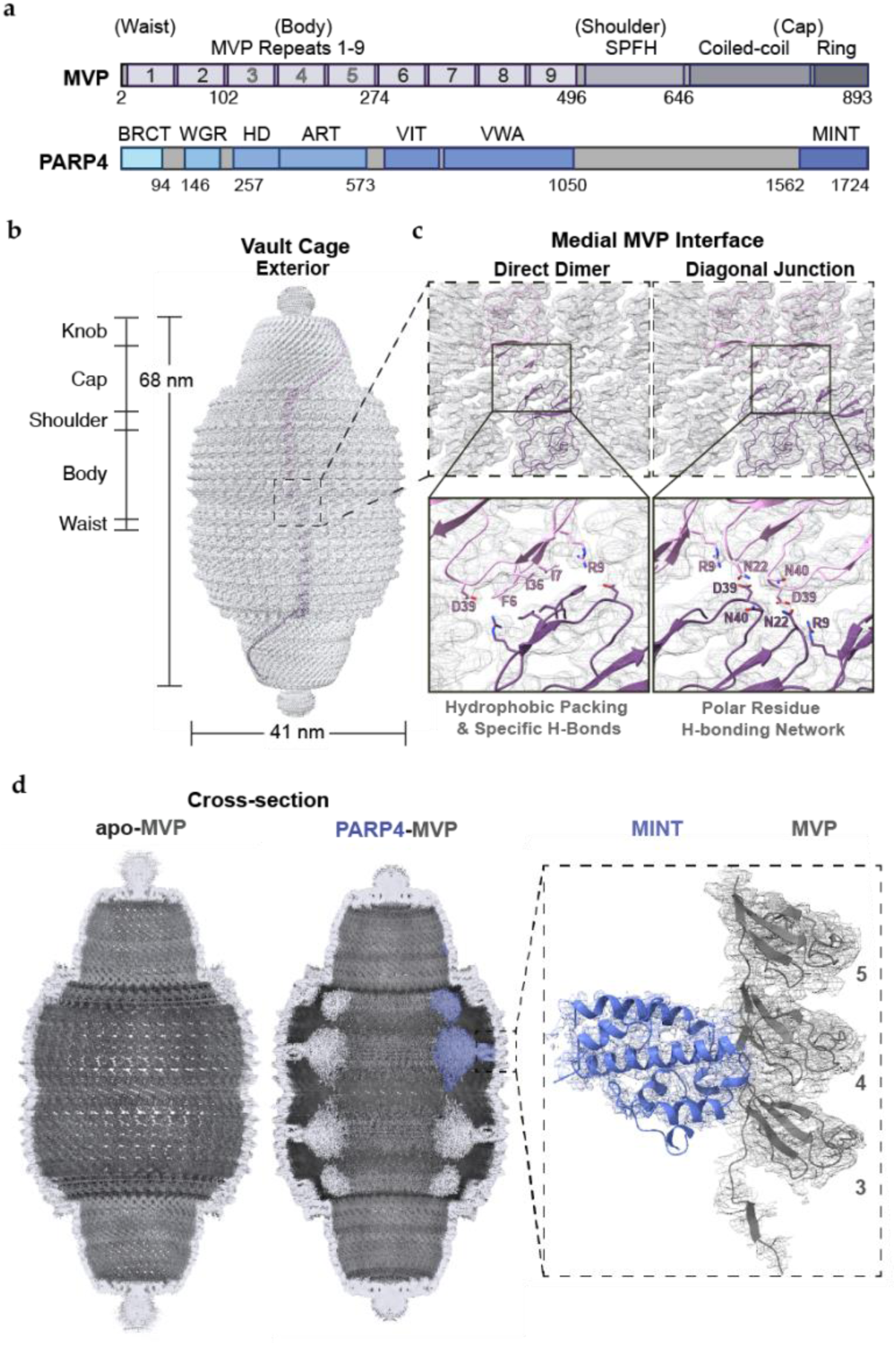
Map of the human vault cage alone and with PARP4. **a**, Domain diagrams of MVP (upper) and PARP4 (lower). **b**, Dimensions and labeled regions of the vault cage, with one MVP monomer from the top half of the cage colored in pink and a medially symmetric monomer from the bottom in purple. **c,** Magnified views of the vault medial interface, with atomic models of MVP fit to the cryo-EM map (gray mesh). Hydrophobic residues F6, I7, and I36 (left), buttressed by two residues with complementary charges (D39 and R9 — center), form the interface between two directly opposed MVP monomers. A network of charged (D39 and R9) and polar (N22, N40) side chains form a dynamic interface between MVP monomers positioned diagonally across opposite halves (right). **d,** Cross-sectional views of the apo- MVP (left) and MVP-PARP4 (right) cryo-EM maps, with potential corresponding to PARP4 colored in blue. Models of the MINT (blue) and MVP3-5 (gray) domains of PARP4 and MVP, respectively, shown within zoned cryo-EM map potential MVP (inset).

## Results

### Structure of the human MVP-PARP4 complex

Full-length constructs of human MVP and PARP4 were cloned into baculoviral vectors for overexpression in insect cells, which do not produce endogenous MVP. An initial attempt to purify the MVP-PARP4 complex by expressing the proteins separately and mixing the cell lysates was unsuccessful, yielding only MVP cages without internalized PARP4 (Supplementary Fig. 1a). Therefore, we changed our expression strategy, simultaneously co-infecting cells with the MVP and PARP4 viruses and adding approximately 5-6 times more PARP4 than MVP virus. This allowed us to compensate for the more robust expression of MVP in insect cells and saturate the PARP4 binding sites within the cage, maximizing the structural information we could obtain about the MVP-PARP4 interface. After the new expression strategy was executed, cell lysate was subjected to exhaustive ultracentrifugation, gradient purification, and gel filtration, from which we recovered pure MVP-PARP4 complexes (Supplementary Figs. 1b and 2a).

Purified human MVP-PARP4, as well as human apo-MVP particles were then applied to carbon-coated grids and subjected to cryo-EM data collection and single particle analysis (Supplementary Figs. 2 and 3). The resulting 3D maps allowed us to compare the human MVP- PARP4 structure to previously published structures of the vault cage. Initially, we performed the reconstruction without enforcing any symmetry and found that the human vault cage adopts the same D39 point group symmetry observed in previously published rat homolog structures [14–16]. Among several differences from these previous structures, however, is the mass of poorly-resolved peptide potential jutting out from the tapered vault “caps.” This peptide signal blurs into something that resembles a “knob" following symmetrization (Fig. 1b). These flexible peptides are likely the product of 30 additional residues at the human MVP C-terminus that are absent in the rat sequence, which is otherwise very similar (sharing 91% sequence identity with human MVP – Supplementary Fig. 4a). Because the knob is close to the symmetry axis and likely to deviate from perfect 39-fold symmetry, it could not be confidently modeled (Supplementary Fig. 4b). However, the presence of the knob suggests that an intact human vault cage is not freely open at the ends.

Our structural model also sheds light on interactions between the symmetrical halves of the human vault, which are thought to separate (either partially or completely) to engulf cargo into the cage [15, 17–19]. Contrary to what was described in the original crystal structure of the rat vault cage, we did not observe salt bridges between E4 and R42 on MVP monomers that are directly opposite one another, despite the fact that the residues were conserved in both species (Supplementary Fig. 4c). This finding was in line with other rat vault solution structures that appeared after the publication of the crystal structure [15, 16], suggesting that the original crystallography conditions may have artificially compressed the cage (Supplementary Fig. 4c-g). In our structure, medially symmetric copies of MVP share a hydrophobic interface to which each monomer contributes three bulky nonpolar residues: F6, I7, and I36 (Fig. 1c). Interestingly, F6 is replaced by an alanine at the same position in the rat MVP sequence (Supplementary Fig. 4c), indicating that the strength of the hydrophobic packing across the halves of the human vault cage may be stronger than that of the rat vault. To either side of the hydrophobic interface, there are two conserved, charged residues, R9 and D39. Although the quality of the map at this relatively flexible region precludes the definitive assignment of their rotamer conformations, R 9 and D39 appear to sit within range to form a salt bridge with one another and form, along with N22 and N40, a loose network of polar residues. These residues are located at the tight junctions between diagonally opposed copies of MVP. Their close proximity and apparent flexibility indicate that these four residues could dynamically interact with each of the others, depending on the local environment or the movement of the cage (Fig. 1c). In addition to contributing to the cumulative strength of the interaction between the cage halves, these polar residues could act as initial points of separation preceding the opening of the vault cage, depending on conditions such as changes in pH, salt concentration, or the introduction of other polar or charged molecules.

### Interface between MVP and PARP4

The most striking feature in our MVP-PARP4 complex structure, however, was the prominent cryo-EM potential inside the vault cage. This additional potential, which corresponded to the PARP4 molecules occupying the cage, was clearly visible in the 3D reconstruction, 2D classes, and even in individual particles (Supplementary Fig. 2) when compared to the particles in the apo-MVP dataset (Supplementary Fig. 3). In a cross-sectional view, the C-terminus of PARP4 can be seen anchored proximal to the vault waist, in between two beta-rich repeat domains of the MVP barrel. It then stretches out into the vault lumen, before the polypeptide chain moves towards the caps, hugging the walls of the cage (Fig. 1d). The long, disordered regions that bridge PARP4’s folded domains, their unconstrained movement in the vault lumen, and the variable occupancy of PARP4 per vault particle across the ensemble, rendered the majority of the enzyme too heterogeneous to be averaged to high resolution. Although processing the data with several different symmetries did not measurably improve resolution for most of the protein structure, imposing D39 symmetry ultimately allowed us to refine the MVP- PARP4 interface to near-atomic resolution (Supplementary Fig. 2). From there, we could dock and refine an AlphaFold2 model of PARP4’s MINT domain within the cryo-EM map (Fig. 1d), making relatively few alterations to the original coordinates (Supplementary Fig. 5a-b). The final refined model revealed the key structural details of PARP4’s recruitment to the MVP cage (Fig. 2a).

**Fig 2.**
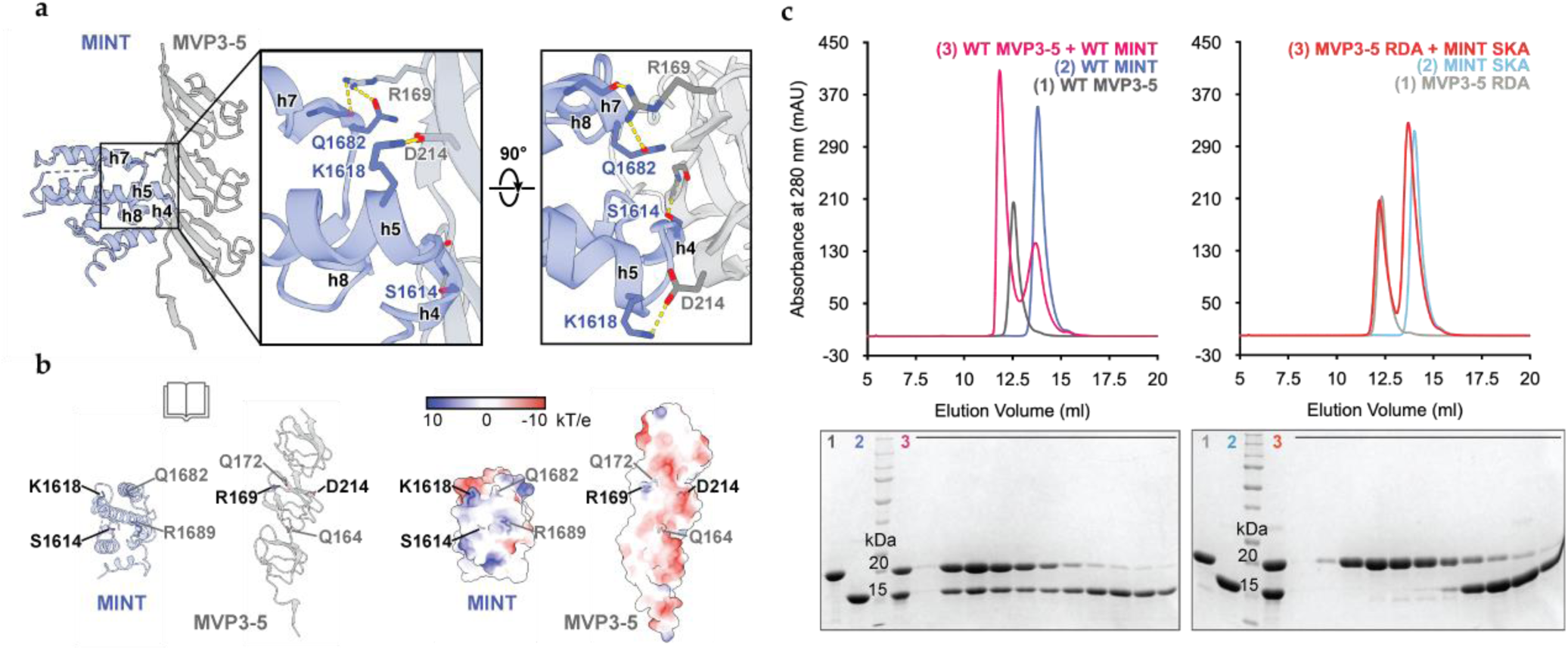
Electrostatic interactions drive MVP-PARP4 complex formation. **a**, Atomic model (left) of the binding interface between the PARP4 MINT (blue) and MVP3-5 repeat domains (gray) from the FL PARP4-MVP structure (inset). MINT domain helices located at the PARP4-MVP interface are labeled with the letter h and associated helix number. Key residues from PARP4 and MVP are labeled in blue and gray, respectively. **b,** The exposed interfaces of MINT (blue) and MVP (gray) are shown in cartoon (left) and surface (right) view representations. Critical interaction residues are labeled in black, with additional residues that may support the interaction shown in gray. Surface views of the MINT and MVP3-5 models are colored by relative electrostatic potential (right), demonstrating electrostatic and steric complementarity at the interface. **c,** Size exclusion chromatograms show individual traces of MVP3-5 (gray) and MINT constructs (blue), indicating their absorbance at 280 nm (upper). Individual traces are overlaid with complex traces following co-incubation of the WT (pink, left) and double mutant constructs (red, right). Corresponding SDS-PAGE gels (lower) from SEC binding assays showing co-elution of WT MVP3-5 and MINT (left) and the separate elutions of double mutant constructs (right) following their respective co-incubations and injections over a Superdex 75 increase column. Black bars denote the area over which fraction samples were collected.

Prior to our structure, only inaccurate models of the MVP-MINT complex generated by predictive softwares existed [20]. Their inaccuracies may have been due to the complications presented by the curvature of the MVP cage with respect to MINT binding since the alignment of bound MINT is out of register with the MVP cage symmetry (Supplementary Fig. 5c). Previously predicted computational models indicated that the MVP-MINT interaction is driven by residues at the very C-terminal end of the MINT domain in its helix-10 [20]. Our structure shows, instead, that residues on MINT’s 4^th^ and 5^th^, as well as its 7^th^ and 8^th^, helices (when starting the count from the N-terminus of the domain) are crucial for MINT’s interaction with MVP (Fig. 2a).

Surface analyses of the MINT-MVP interface reveal that it falls in a region of substantial electrostatic complementarity between MVP and PARP4. Specifically, an acidic stretch that runs along MVP’s third and fourth repeat domains aligns with a basic region between helices-4 and - 5 of the MINT domain. Within this region, PARP4’s K1618 and MVP’s D214 residues forge a salt bridge, while several nearby residues engage in additional, specific electrostatic interactions. Notably, the side chains of PARP4 S1614 and R1689, as well as MVP R169 and Q164, appear to hydrogen bond with backbone amide groups of the opposite species. The side chains of PARP4 Q1682 and MVP Q172 are in range to hydrogen bond with one another as well (Fig. 2b). Having identified these residues as important drivers of complex formation, we sought to validate our structural results by mutating several of them to alanines and measuring the impact on binding.

We designed PARP4 and MVP constructs that contained only the necessary interaction domains (MINT_1562-1724_ and MVP3-5_102-274_, respectively), both of which could be readily overexpressed in and purified from *E. coli*. Prior to injecting them over a gel filtration column, we incubated approximately equimolar concentrations of wildtype (WT) MINT and WT MVP3-5. The resulting peak eluted at a lower volume than the elution volumes of each individual species. We confirmed that they comigrated by SDS-PAGE gel (Fig. 2c). When we pooled fractions from the complex peak and re-injected the sample over the column, we observed a peak that eluted at the same volume, demonstrating the considerable stability of the complex (Supplementary Fig. 6a). After we confirmed that these domains could bind and remain stably intact over a size exclusion column, we mutated some of the previously identified interacting residues (the PARP4 residues S1614 and K1618 and MVP residues R169 and D214) to alanines. We dubbed the respective constructs the “SKA” and “RDA” mutants and used gel filtration to assay each construct’s ability to bind the corresponding species. Both of the mutant constructs (Fig. 2c), as well as combinations of each mutant and WT species (Supplementary Fig. 6b-c), were co-incubated prior to injection over a gel filtration column. In each case, we saw no evidence of complex formation, indicating that the interaction between PARP4 and MVP relies on specific electrostatic interactions between key residues.

### Structure of the NAD^+^-Bound MVP Cage

As a complement to our structure of MVP bound to PARP4 alone, we sought to collect a cryo-EM dataset of the complex in the presence of NAD^+^ to determine whether active PARP4 would adopt an alternate conformation or whether its activity would influence the conformation or stability of the vault cage. Our previous observation that polar and charged residues stabilized inter-half MVP interactions led us to hypothesize that PARP4 could modify these residues with bulky ADP-ribose groups, consequently triggering an initial separation between the halves that could destabilize the cage and permit cargo entry. To test whether active PARP4 would destabilize the cage, we incubated the MVP-PARP4 complex with 1mM of NAD^+^ for several hours, dialyzed excess NAD^+^ out overnight, and froze grids with the complex solution for data collection. Contrary to our expectations, introducing NAD^+^ did not appear to decrease the proportion of intact vault cages on the grid. However, after collecting and processing a full dataset of these MVP-PARP4 complexes in the presence of NAD^+^, we noticed some additional, previously unobserved cryo-EM potential within the interior of the SPFH domains in the “shoulder” region of the vault cage. We soon found that we could dock an NAD^+^ molecule into this potential (Fig. 3a-b). Notably, we also observed changes in MVP’s structure near the binding site, providing additional evidence for the notion that MVP binds NAD^+^ directly. Namely, when NAD^+^ was present, several residues in the binding site adopted different rotamer conformations, and a loop appeared to be displaced from its previous position directly in front of the pocket (Fig. 3b). This newly discovered ligand binding site sits within the vault cage interior at the interface between the SPFH domains of two MVP monomers (“MVP1” and “MVP2)”, nestled behind the “keyhole loop,” which extends into the vault lumen and creates a small pocket between itself and the remainder of the domain (Fig. 3c).

**Fig 3.**
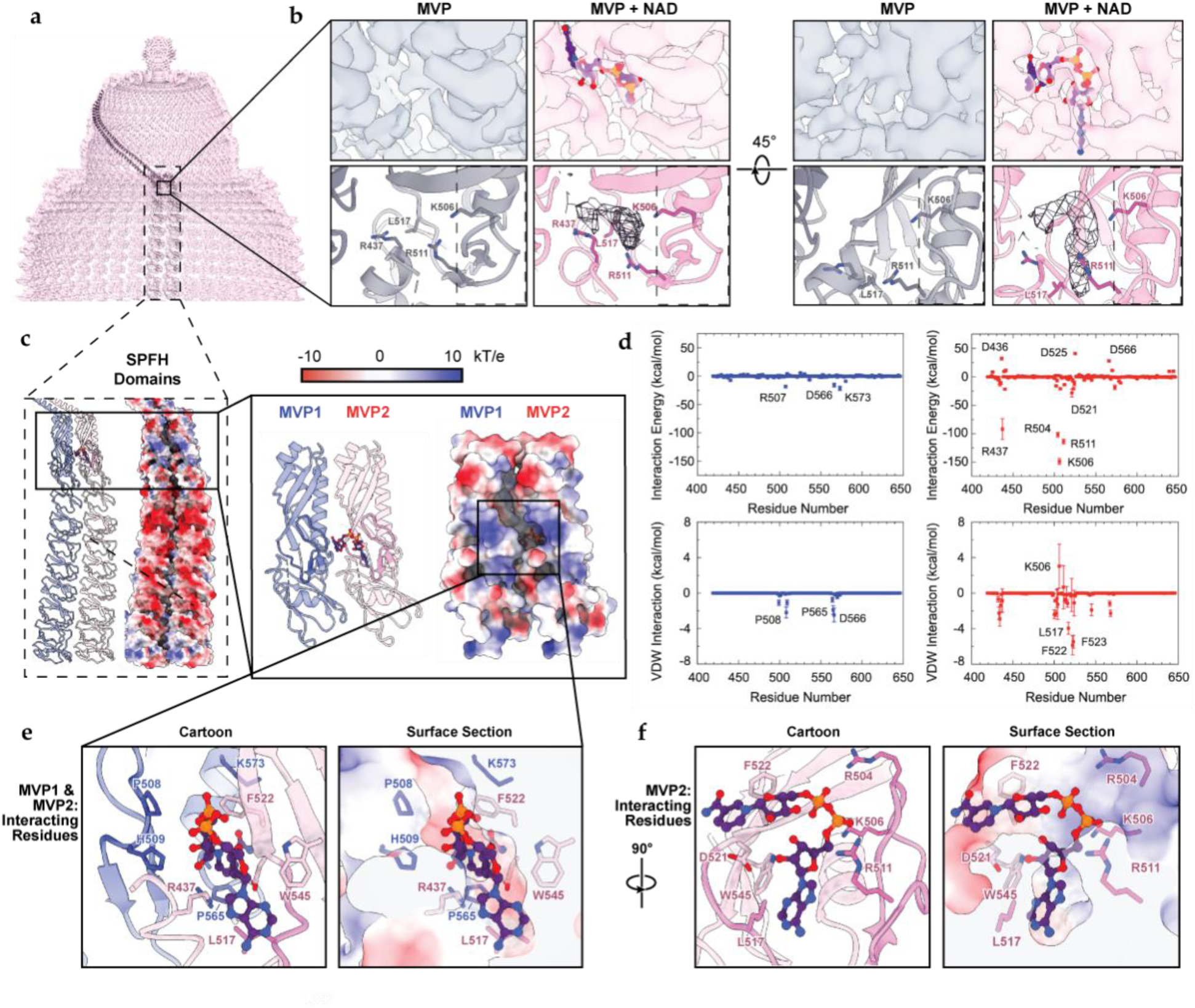
NAD^+^ Potential Behind the MVP Keyhole Loop. **a**, Cross-section of the upper half of the MVP cryo-EM map, with adjacent MVP monomers (black, cartoon view) fit into the potential. **b**, Magnified top view (left) of the hooked ligand signal in the MVP-PARP4-NAD^+^ map (pink) compared to the corresponding vacant site in the MVP-PARP4 map (gray). An atomic model of NAD^+^ is docked into each site at equivalent positions to highlight the absence of ligand signal in the MVP-PARP4 map (lower panels), with residues that adopt different rotamers labeled in gray and pink, respectively. Front-facing views of the same maps, focused on MVP’s NAD^+^-binding site (right). **c,** Adjacent NAD^+^-bound MVP monomers MVP1 and MVP2 in cartoon view (colored blue and pink respectively, left) and in surface view (colored by electrostatic potential, right). Magnified views of MVP1 and MVP2 SPFH domains, with the keyhole loop shown in darker blue and pink respectively (inset, left). Surface view of the same model (inset, right). Bound NAD^+^ is shown in purple. **d,** Pairwise interaction energies between individual MVP residues and NAD^+^ moieties calculated by MD simulations (right). Results corresponding to residues from MVP1 are shown in blue, and those for MVP2 are shown in red. **e,** Atomic model of MVP’s NAD^+^-binding site. Side chains (stick view) of residues from MVP1 (blue) and MVP2 (pink) that comprise the boundaries of the site are shown and labeled in cartoon (left) and electrostatic surface view (right). Cryo-EM map signal is shown as black mesh, and an atomic model of NAD^+^ is docked into each site at equivalent positions to show ligand position with respect to the relevant residues. **f,** Magnified view of the NAD^+^- binding site, with crucial MVP2-interacting residue side chains labeled in pink.

To verify that the potential represented bound NAD^+^ and not covalently attached ADP-ribose, we employed two experimental strategies. First, we obtained the structure of MVP alone (without the ADP-ribosyltransferase PARP4) in the presence of NAD^+^, and then we conducted a mass spectrometry-based analysis to detect ADP-ribosylation on the vault cage. In our MVP- NAD^+^ cryo-EM map, we observed the same ligand potential in the SPFH domain pocket, indicating that MVP specifically binds NAD^+^ and that PARP4 is not necessary for the interaction (Supplementary Fig. 7a). Further, after incubating NAD^+^ with both the MVP-PARP4 complex and MVP alone and submitting both samples for LC-MS/MS, we were unable to identify an ADPr modification near the binding pocket by LC-MS/MS in either sample. This provided additional evidence that the cryo-EM potential corresponded to non-covalently bound NAD^+^.

Whereas the majority of vault’s interior surface is relatively smooth and acidic, the binding site of NAD^+^ is a recessed pocket lined by basic residues (Fig. 3c), which we determined to be crucial for recruiting and stabilizing NAD^+^.

### Biochemical Basis for the MVP-NAD^+^ Interaction

In order to better understand this interaction, we performed molecular dynamics simulations to calculate pairwise interaction energies between MVP side chains and NAD^+^ moieties. The results suggested that interaction energies between NAD^+^ and basic residues from MVP2 (R504, K506, and R511) and NAD^+^ are extremely favorable and likely drive the interaction via salt bridge formation (Fig. 3d). However, the copy of MVP that sits to the left of NAD^+^ (MVP1) also supports the ligand’s association to the vault cage, largely by applying additional steric constraints to the binding site and contributing to the Van Der Waal (VDW) interactions. In particular, P508, H509, and P566 help to define the left side boundary of the pocket, while H509’s side chain also appears to hydrogen bond with the 2’-OH of the distal ribose group of NAD^+^. MVP1’s R507 and K537 residues appear to further neutralize the negative charge introduced by NAD^+^’s diphosphate group (Fig. 3e). In addition to the aforementioned basic residues on MVP2’s keyhole loop (R504, K506, and R511), the residues R437, D521, and W545 hydrogen bond with the ribose moieties of NAD^+^. Three hydrophobic residues from MVP2, L517, F522, and F523, serve to delineate the boundaries of the pocket, with L517’s side chain rotamer position also changing in the presence of nucleotide (Fig. 3f).

Together, our structural and simulation data depict a well-defined, novel NAD^+^-binding pocket between the SPFH domains of two adjacent MVP monomers, whose electrostatic properties are ideal for stabilizing NAD^+^’s diphosphate and ribose groups at the site.

Additionally, we performed isothermal titration calorimetry (ITC), which suggested that NAD^+^’s association to the vault cage exists in the micromolar range (Supplementary Fig. 7b). However, this estimation may not be accurate since the intact vault cage is not highly permeable to metabolites of a size comparable to NAD^+^. Therefore, the time allowed between ITC injections may be insufficient for most of the available NAD^+^ to enter the cage.

To explore the potential versatility of the ligand binding site, we conducted additional MD simulations in which ligands structurally akin to NAD^+^ were positioned at the binding site, with each simulation running for 2 µs. While adenosine diphosphate (ADP) and guanosine diphosphate (GDP) could both stably occupy the pocket for that length of time, an adenine base quickly escaped the site (Supplementary Fig. 7c). These results indicated that a purine base together with a diphosphate group are likely adequate for a ligand’s association to the vault cage. To further investigate the potential interaction, we determined the cryo-EM structure of human MVP in the presence of ADP. Similar to the NAD^+^-bound structure, the cryo-EM map exhibited additional potential behind the keyhole loop. However, this potential was not as distinct or as prominent as that found in the MVP-NAD^+^ map, nor was the potential for rotamers of key sidechains around the ligand binding site as distinct (Supplementary Fig. 8). This indicated that ADP occupancy was likely lower and therefore more likely to be averaged out.

The lower apparent occupancy of ADP in the vault cage suggests that, while the purine diphosphate groups are likely adequate for MVP binding, NAD^+^’s additional ribose moieties strengthen the interaction.

Identifying the NAD^+^-binding site prompted us to ask if there was a functional link between vault’s resident PARP enzyme and MVP’s ability to harbor its enzymatic substrate (given the importance of NAD^+^ availability in PARP regulation [21]). Intriguingly, the volume of the PARP4 potential adjacent to MVP’s SPFH domain can accommodate PARP4’s catalytic region (comprising its HD and ART domains), inviting the possibility that the vault cage may act as an NAD^+^ reservoir for vault-bound PARP4 (Fig. 4a-b). To further probe the relationship between MVP’s NAD^+^- and PARP4-binding sites, we analyzed the conservation of key residues at those sites by performing a multiple sequence alignment (MSA) of MVP from several representative vertebrate and invertebrate species. We found that the conservation patterns of MVP residues involved in PARP4 binding (present only in the vertebrate species) differed from those responsible for the NAD^+^ interaction (which had variable conservation between the two groups) (Fig. 4c). This could reflect a model in which PARP4 began to opportunistically occupy the vault in higher order organisms to take advantage of the stored NAD^+^ as an enzymatic substrate.

**Fig 4.**
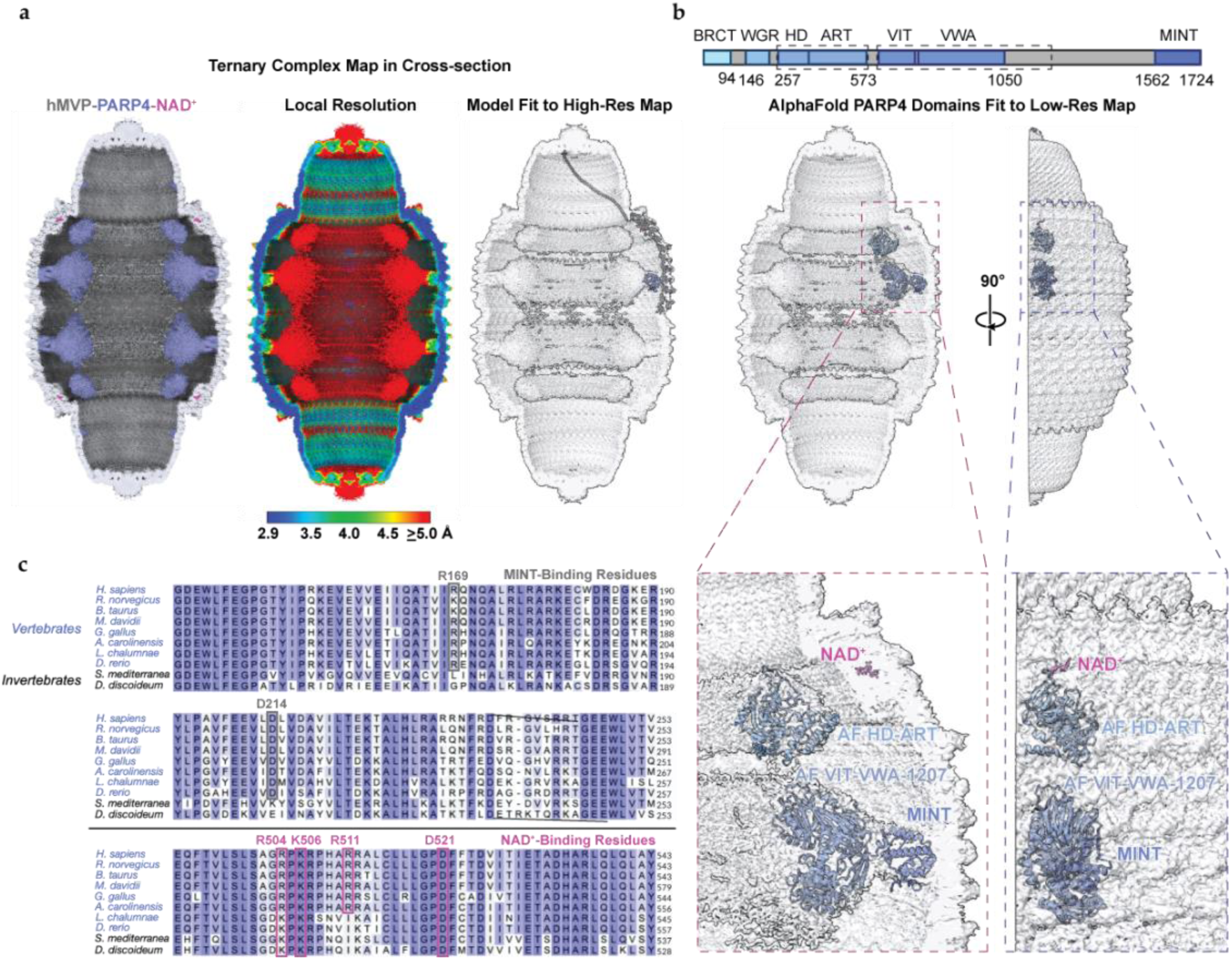
Human PARP4 may opportunistically consume MVP-bound NAD^+^. **a**, Cross-sectional views of the MVP-PARP4-NAD^+^ (ternary complex) cryo-EM maps, with color coding by molecular species (left) and local resolution (center), as well as a transparent view of the map with the MVP-MINT-NAD^+^ molecular model docked into the high-resolution region (right). **b,** Domain diagram of PARP4 (upper) with the HD-ART and VIT-VWA regions highlighted by dashed line boxes. Cross-sectional view of the ternary complex map with AlphaFold2 models of the boxed PARP4 domains docked into it, along with MINT and NAD^+^ for reference (left). View of the docked models turned 90° to show their alignment in the map (right). Magnified views of the docked models (insets, lower). **c,** Multiple sequence alignment (MSA) of the PARP4 and NAD^+^-binding regions of MVP in a representative sample of vertebrate (blue) and invertebrate (gray) species. Residues most critical for human MVP’s interaction with PARP4 and NAD^+^ are outlined in grey and pink, respectively.

### Vault coimmunoprecipitation (co-IP) from WT and PARP4-depleted cells

In order to find native vault interaction proteins, we developed an anti-human MVP synthetic antigen binding reagent (sAB) that could be biotinylated and subsequently used for proteomic analysis (Fig. 5a). We chose to use A549 cells (derived from human lung epithelia) as our native vault source due to their high constitutive expression of vault proteins and vault’s documented, medically relevant activity in lung tissue [2, 3, 22]. We engineered a PARP4- depleted cell line using CRISPR/Cas9 so that we could determine PARP4’s influence on the vault interactome. We verified PARP4 depletion via sequencing and Western blot (Fig. 5b).

**Fig 5.**
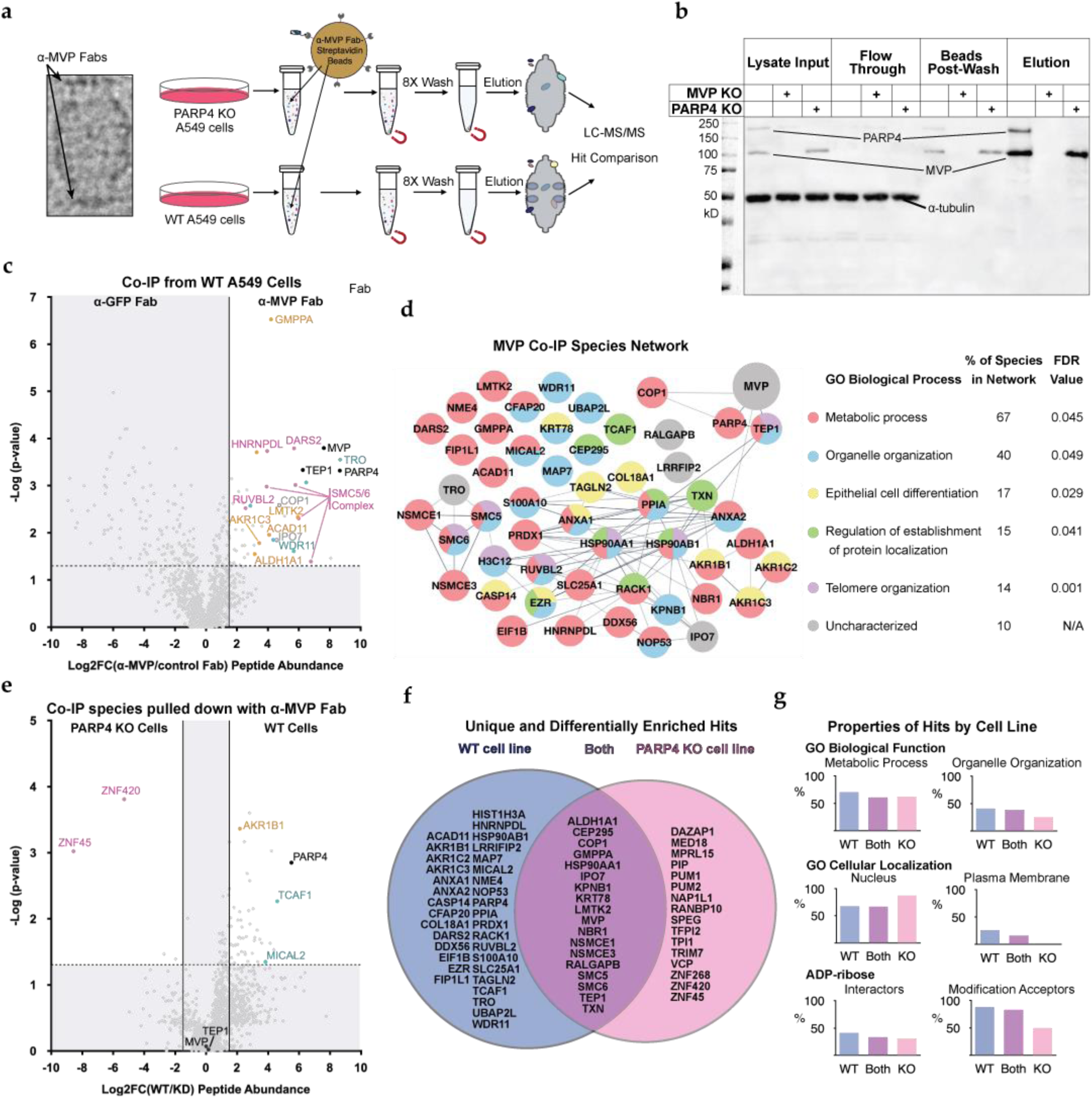
Depleting PARP4 alters the vault interactome. **a**, Cryo-EM micrograph image of the MVP cage with a ring of α-human MVP Fabs (indicated by arrows) bound to each cap (left). Schematic of anti-MVP coimmunoprecipitation experiment (right). **b,** anti-MVP and -PARP4 western blot demonstrating the anti-MVP sAB’s ability to pull down vault proteins from WT and PARP4 KO A549 cells. The whole lysate input, flow through, residual protein bound to streptavidin beads, and elution fractions from WT, MVP KO, and PARP4 KO cells were run on an SDS-PAGE gel following MVP coimmunoprecipitation (co-IP) and transferred to a PVDF blot. Proteins were incubated with species-specific rabbit primary anti-bodies, then anti-rabbit IgG-HRP secondary antibodies for detection with chemiluminescent HRP substrate. α-tubulin was imaged as a loading control. **c,** Volcano plot showing differential enrichment of species co-IP’ed with the anti-MVP (right) and anti-GFP control (left) sABs from WT A549 cells. Thresholds for “hits” (log2 fold change of peptide abundance from the experimental versus control IP condition > 1.5 and significance of value across triplicate experiments > 1.3) shown as dashed lines. Relevant enriched hits are labeled, with vault components written in black text. **d,** Network of hits co-IP’ed with MVP from WT A549 cells, with connecting lines representing previously established interactions between the proteins (left). Key showing significant gene ontology (GO) biological process terms associated with the network, the proportion of proteins tagged with those terms, and their false discovery rates (FDR) (right). **e,** Volcano plot showing differential enrichment of species co-IP’ed with the anti-MVP sAB from WT A549 (right) and PARP4 KO A549 cells (left). Thresholds for WT vault “hits” shown as dashed lines. Relevant enriched hits are labeled, with vault components written in black text. **f,** Venn diagram showing hits that were either unique to or significantly enriched in the WT (versus PARP4 KO) cell co-IP dataset (blue, left), hits that were unique to the PARP4 KO cell co-IP dataset (pink, right), or hits present in both datasets without significant differential enrichment in either (purple, center). **g,** Bar graphs depicting the proportion of proteins in each dataset tagged with specific GO biological process (upper) and GO cellular component terms (center), as well as the proportion of those previously observed to either interact with (left) or be modified with (right) ADP-ribose (lower).

Significant hits from our co-IPs (with log_2_(Fold Change) ≥ 1.5 and -log_10_(p-value) > 1.3) included known MVCs (black data points, Fig. 5) and COP1 (an E3 ubiquitin ligase previously observed to interact with vault [23]), which validated our approach. We identified a total of 52 hits from the WT A549 cells (Fig. 5c) and subjected them to gene ontology (GO) network analysis (Fig. 5d) to evaluate their biological roles and cellular compartmentalization. We then compared the co-IP hits from the WT and PARP4-depleted cell lines, sorting them into three groups. 34 “WT hits” (Fig. 5e) were either exclusively identified or significantly enriched in WT cells, while 16 “KO hits” were uniquely present in our PARP4-depleted cells. 18 “shared hits” were identified in both datasets without significant differential enrichment in either (Fig. 5f).

Many of the most overrepresented genes in all three groups were active in pro-survival and pro-proliferative pathways, including a number of cytosolic enzymes involved in nucleotide metabolism (gold data points, Fig. 5) and proteins that operate at the gene regulation level (pink data points, Fig. 5). WT hits in the “gold set” included ACAD11, AKR1C3, and AKR1B1, all nucleotide-binding dehydrogenases that promote cell growth and survival in the wake of stress [24, 25], while those in the “pink set” included RUVBL2 [26], HNRNPDL [27], and DARS2.

Although it is canonically known as a tRNA synthetase, DARS2 has recently been implicated in maintaining mitochondrial metabolism by regulating splicing [28] and transcription [29]. A third set of highly abundant genes in our pulldowns regulate cell migration and invasion (teal data points, Fig. 5), including TRO [30], one of the most abundant hits, as well as RACK1 [31], WDR11 [32], MICAL2 [33], and — the only gene in this group to inhibit cell migration — TCAF1 [34]. Finally, SLC25A1, a solute carrier, emerged as an interesting hit for its homology with another solute carrier that transports NAD^+^ into the mitochondria [35]. Broadly speaking, most WT hits were associated with either the “metabolic process” or “organelle organization” GO biological process terms. Their “cellular compartment” GO terms (derived from the Ensembl database) place them throughout the cell, with a little over a quarter able to localize to the plasma membrane and a little over half present in the nucleus.

By contrast, according to their cellular compartment GO terms, none of the KO hits localize to the plasma membrane, and over three quarters spend time in the nucleus. Further, fewer KO hits were associated with organelle organization than WT hits, though a similar proportion were involved in metabolic processes (Fig. 5g, Supplementary Table 1). Like the WT hits, many of the KO hits belonged in the “pink set” for their ability to regulate cell growth and survival at the gene expression level. Among them were the KRAB-type zinc fingers ZNF420, a known negative regulator of apoptosis [36], and ZNF45, whose expression has been observed to be upregulated in some cancer models [37] [38]. Three functionally related RNA-binding proteins also emerged as prominent hits, all of which have been shown to suppress gene expression following interactions with the 3’UTRs of various mRNAs [35]. The most abundant of these, DAZAP1, has also been implicated in splicing events that promote cell growth [39], while the others (PUM1 and PUM2) have been associated with cancer cell proliferation [40].

The shared hits exhibited GO functions and subcellular localization patterns broadly similar to those of the WT hits (Fig. 5g). “Gold set” pro-growth/-survival metabolic enzymes include LMTK2 and ALDH1A1 (which uses an NAD cofactor) [41], while “pink set” hits that regulate gene expression were represented by components of the Smc5/6 complex. This assembly is a cohesin/condensin-like structure that promotes cell growth and protects the cell from genotoxic stress (and even viral infection) by interacting with DNA and mediating homologous recombination (HR) [42, 43]. The complex’s ability to regulate HR at telomeres also links SMC5 and SMC6 to organelle organization and “telomere organization” functions in the GO network analysis, a function shared by several other hits, including the MVC TEP1. Hits with the “organelle organization” designation also included KPNB1, which, along with IPO7, regulates nucleocytoplasmic shuttling [44], a previously hypothesized function for vault. Finally, GMPPA, which mediates protein glycosylation as a regulatory subunit in the GDP-mannose biosynthetic enzyme complex, has been linked to tumor growth [45] and chemotherapeutic drug resistance [46], pathologies previously linked to MVP upregulation [4].

## Discussion

Our structure of the MVP-PARP4 complex resolves a longstanding question in the field regarding the biochemical basis for PARP4’s recruitment to the vault cage. Along with shedding light on vault’s native function, this structure could inform efforts to develop vault- based biomolecule delivery systems that use MINT or mutants thereof as a fusion domain [13]. In addition, our discovery of MVP’s novel NAD^+^-binding pocket introduces several new questions to the field. Provided that NAD^+^ is MVP’s preferred ligand, its association to the vault cage could have a number of functional consequences. By analogy to the functions of other non-enzymatic NAD^+^-binding proteins [47], MVP’s interaction with NAD^+^ could potentially stabilize a particular conformational state [48, 49] (thereby altering MVP’s binding properties [50]) or influence NAD^+^ compartmentalization and availability for enzymes that require it [35]. Both functions appear possible. With respect to the former, bound NAD^+^ displaces a loop just beneath MVP’s SPFH domain, appearing to push it further into the lumen (Fig. 3b) where it could potentially interact with vault cargo or MVCs more readily. As to the latter function, NAD^+^’s proximity to PARP4, as well as our discovery of vault cargo molecules that bind NAD (Supplementary Table 1), could signal vault’s ability to act as a repository for enzymes that utilize NAD^+^ as a co-factor or substrate. In either case, MVP’s capacity to bind NAD^+^ indicates that vault may be able to adapt its behavior in response to changes in cellular metabolite concentration, offering one potential explanation for its functional versatility in different cell types and conditions.

PARP4’s role in the vault complex still requires investigation. However, our proteomics data indicate that vaults from WT and PARP4-depleted cells exhibit distinct proteins in their interactomes, suggesting that PARP4 may play a role in regulating vault’s cargo selection or subcellular localization. “KO hits” from PARP4-depleted cells were more likely to be found in the nucleus. Accordingly, they were often associated with gene regulation and nucleic acid binding. By contrast, many “WT” hits localized to the plasma membrane and regulated cell migration and invasion functions. PARP4’s VWA domain (a protein-protein interaction domain common in cell surface proteins that regulate cell migration [51]) could conceivably facilitate vault’s recruitment of these cargo proteins. We have not yet determined whether PARP4’s catalytic activity is influencing vault cargo recruitment or if the cargo molecules themselves are being modified. Our analysis of WT and KO hits indicates that a similar proportion of genes in each dataset have previously been observed to bind ADP-ribose. [52, 53] However, a greater proportion of WT hits than KO hits have been observed to accept ADP-ribose modifications [54]. This subset of WT hits could constitute some of vault-bound PARP4’s modification targets (Fig. 5g, Supplementary Table 1). Further exploration of these molecules could thus shed light on vault’s roles in cell signaling. More broadly, our identification of native vault-binding partners involved in cell growth, proliferation, chemoresistance, and migration can begin to explain MVP’s previously observed roles in cancer progression [55, 56] and drug resistance [4], as well as metabolic regulation [7], a function now further cemented by our discovery of its association with the critical metabolite NAD^+^.

## Methods

### Plasmid Construction and Baculovirus Generation

Human MVP and PARP4 genes were synthesized by GenScript and subcloned into the pVL1393 baculovirus transfer vector. pVL1393-hMVP and pVL-hPARP4 plasmids were incubated with BestBac2.0 linearized baculoviral DNA (Expression Systems, 91-002) and transfected into adherent Sf9 insect cells using Cellfectin II (Invitrogen 10362-100). After 7 days, the cells were pelleted by centrifugation and P0 baculovirus was harvested from the supernatant fraction. P0 virus was amplified in Sf9 suspension cells for an additional week to generate P1 baculovirus stocks. P1 viruses were, in turn, amplified in Sf9 cells to generate P2 stocks. P1 and P2 stocks were used to infect Hi5 cells for protein expression. Constructs for *E. coli* expression of WT and mutant MVP3-5 and MINT domains were cloned into the pET47b(+) vector, which introduced an N-terminal His-tag and HRV 3C cut sites. Alanine mutations were imposed by site-directed mutagenesis and confirmed by Sanger sequencing.

### Expression and Purification of Vault Cage Complexes

Hi5 cells were grown to one liter at a density of about 2 x 10^6^ cells/ml and infected with 1 ml hMVP P1 baculovirus. Cells grown to produce the MVP-PARP4 complex were simultaneously infected with 5-6 ml hPARP4 P2 baculovirus. Infected cells were rotated on an orbital shaker at 120 rpm at 27°C. After 65 hours, they were harvested by centrifugation at 900 *g* for 15 minutes. Cell pellets were washed with 20 ml of phosphate buffered saline (PBS) per 0.5 L of cells, transferred to 50 ml Falcon tubes, and centrifuged for an additional 15 minutes at 900 *g*. Cells were decanted, and aliquots were flash frozen in liquid nitrogen prior to long term storage at -80°C. For purification, 0.5 L cell aliquots were thawed in room temperature water for 10 minutes, then resuspended in 120 ml Buffer A (50 mM Tris pH 7.4, 75 mM NaCl, 1.5 mM MgCl_2_, & 1 mM DTT), with 1% Triton X-100 and 1 mM phenylmethylsulfonyl fluoride (PMSF).

Pellets were disrupted with 30-40 strokes of a Type A Dounce Homogenizer. Lysate was transferred to centrifuge tubes and vortexed twice during a twenty-minute incubation on ice. Lysate was cleared by centrifugation at 20,000 *g* for 15 minutes, and the supernatant was harvested for additional purification steps. High-molecular weight complexes were pelleted by centrifugation at 150,000 *g* for one hour. Pellets were resuspended in 6 ml of Buffer A with 7% Ficoll and sucrose added. The resuspension was centrifuged for 45 minutes at 43,000 *g* in order to pellet microsomal contaminants. The supernatant fraction was diluted in 21 ml of Buffer A, and pelleted by ultracentrifugation for 3 hours at 200,000*xg*. Pellets were resuspended in 0.8 ml of Buffer A and incubated with 500 µg RNase A as well as 50 U of RNase T1 for 15 minutes.

Denatured ribosomal proteins were pelleted by centrifugation at 30,000 *g* for 20 minutes. The supernatant was then transferred to a new 1.5 ml tube for an additional 15 minute spin. The remaining supernatant was taken up to 1 ml with Buffer A and gently pipetted over a 20%-60% sucrose gradient. Gradients were centrifuged at 78,000 *g* for 16 hours. Fractions from the 45% and upper half of the 50% gradient fractions were pooled and dialyzed in ADP-ribosylation buffer (50 mM HEPES pH 8, 5 mM MgCl_2_, 5 mM CaCl_2_, 0.25 mM DTT) overnight. Dialysate was concentrated to 500 µl and injected over a Superose 6 increase 10/300 GL column (Cytiva). Fractions were aliquoted and flash frozen with liquid nitrogen prior to storage at -80°C.

### Expression and Purification of MVP3-5 and MINT Truncation Constructs

pET47b-MVP3-5 and pET47b-MINT WT and mutant constructs were transformed into Rosetta Singles BL21(DE3) *E. coli* competent cells. Protein production was induced by the addition of 0.5 mM isopropyl β-D-1-thiogalactopyranoside (IPTG) at OD_600_ 0.6-0.75, and cells were kept at 18°C and rotated at 220 rpm overnight. For purification, cells were pelleted by centrifugation at 6,000 rpm for 20 minutes and each pellet was resuspended in 50 ml Ni-NTA buffer (20 mM Na_2_HPO_4_/NaH_2_PO_4_ pH 7.4, 300 mM NaCl, 1 mM PMSF) with 10 mM of imidazole. Resuspended cell solutions were subjected to sonication for 3 minutes at the 75% amplitude setting, with a pulse sequence of 3s on/5s off. Lysate was cleared by centrifugation at 16,500 rpm for 45 minutes, and the supernatant was flowed over 6 ml of washed Ni-NTA resin (ThermoFisher Scientific) in a gravity column. Resin beads were washed with 150-200 ml Ni-NTA buffer with 20 mM Imidazole. Proteins were eluted from the column following five minute incubations in 15-25 ml Ni-NTA buffer with 250 mM Imidazole. Protein eluate was dialyzed into 2 L SEC buffer (50 mM Tris pH 8, 150 mM NaCl, and 0.5 mM tris(2-carboxyethyl)phosphine (TCEP-HCl)) and subjected to His-tag cleavage with PreScission Protease overnight. Dialyzed protein was concentrated, filtered and injected over a Superdex 75 increase 10/300 GL column (Cytiva). Peak fractions were pooled and used immediately for gel filtration peak shift assays.

### Cryo-electron Microscopy Sample Preparation

Vault cage peak fractions were concentrated and incubated at 4°C for 3.5-4 hours in the presence 1 mM NAD^+^ or ADP or in the absence of ligand. Samples were dialyzed in ADPr reaction buffer overnight to remove excess metabolite in the ligand-added solutions. Vault proteins were then concentrated and subjected to 5 minutes of centrifugation at 13,000 rpm to clear aggregates. Quantifoil (R1.2/1.3, 200 mesh) copper grids with 2 nanometers of continuous carbon film (Electron Microscopy Sciences) were glow discharged using a 15 second, 5 watt program on the Solarus 950 Plasma Cleaner System (Gatan). Three microliters of sample (∼1.5-1.8 mg ml^-1^) were applied to the carbon-coated grid surface. The climate chamber of the Vitrobot Mark IV (ThermoFisher) was set to 8°C and 100% humidity. Grids were blotted for 1-2 seconds at blot force 0 with standard Vitrobot filter paper (Ted Pella, 47000-100), then plunge-frozen into liquid ethane.

### Cryo-electron Microscopy Data Collection

Data were collected at one of two facilities, both of which used a 300 kV Titan Krios transmission electron microscope (ThermoFisher Scientific). At the Advanced Electron Microscopy Facility at the University of Chicago, movies were recorded on a K3 direct detector camera (Gatan) at a nominal magnification of 64,000x (translating to pixel size of 0.672 Å at the detector), in super-resolution counting mode by image shift. EPU software was set to automated acquisition mode and collected (depending on the dataset) between 2745 and 6968 image stacks, each with 40 frames, subject to a total dose of 45 e^-^/Å^2^. The defocus range was set to -1.0 to -2.5 μm. At the HHMI Janelia Farms Cryo-Electron Microscopy Facility, movies were recorded on a K3 Bioquantum camera (Gatan) at a nominal magnification of 53,000x (translating to a pixel size of 0.66 Å at the detector), in super-resolution counting mode by image shift. SerialEM software was used to collect data using a strategy of 3 by 3 + 1 shots per hole, acquiring (depending on the dataset) between 3692 and 7491 image stacks. See **Table 1** for additional details.

**Table 1.** Data Collection and Refinement Statistics.

|  | Human MVP | Human MVP-NAD<sup>+</sup> | Human PARP4-MVP | Human PARP4-MVP-NAD<sup>+</sup> | Human MVP-ADP | Human MINT-MVP |
| --- | --- | --- | --- | --- | --- | --- |
| **Data collection and processing** |  |  |  |  |  |  |
| Magnification | 64,000 | 64,000 | 53,000 | 53,000 | 53,000 | 64,000 |
| Voltage (kV) | 300 | 300 | 300 | 300 | 300 | 300 |
| Electron exposure (e-/Å<sup>2</sup>) | 45 | 45 | 45 | 45 | 45 | 50 |
| Defocus range (µm) | -1.0 to -2.5 | -1.0 to -2.5 | -0.7 to -1.7 | -0.7 to -1.7 | -0.7 to -1.7 | -1.0 to -2.5 |
| Pixel size (Å) | 1.344 | 1.344 | 1.32 | 1.32 | 1.32 | 1.344 |
| Symmetry imposed | D39 | D39 | D39 | D39 | D39 | D39 |
| Initial particle images (no.) | 205,194 | 318,989 | 670,079 | 174,624 | 214,820 | 96,257 |
| Final particle images (no.) | 98,674 | 108,991 | 44,443 | 93,859 | 100,860 | 19,104 |
| Map resolution (Å) | 3.31 | 3.07 | 2.92 | 2.93 | 2.68 | 3.40 |
| FSC threshold | 0.143 | 0.143 | 0.143 | 0.143 | 0.143 | 0.143 |
| **Refinement** |  |  |  |  |  |  |
| Initial model used (PDB code) | 46V0 |  | 46V0 | 46V0 |  |  |
| Model resolution (Å) | 3.64 |  | 3.47 | 3.49 |  |  |
| FSC threshold | 0.5 |  | 0.5 | 0.5 |  |  |
| Map sharpening B factor (Å<sup>2</sup>) | -124.7 | -120.1 | -81.5 | -80.5 | -104.0 | -87.9 |
| **Deposition Accession Codes** |  |  |  |  |  |  |
| EMD | 44953 | 44954 | 44955 | 44957 | 44959 | 44960 |
| PDB | 9BW5 |  | 9BW6 | 9BW7 |  |  |
| **Model composition** |  |  |  |  |  |  |
| Nonhydrogen atoms | 12,300 |  | 14,604 | 14,692 |  |  |
| Residues | Protein: 1,550 |  | Protein: 1,838 | Protein: 1,838 |  |  |
| Ligands | — |  | — | NAD<sup>+</sup>: 2 |  |  |
| **B factors (Å<sup>2</sup>)** |  |  |  |  |  |  |
| Protein | 83.01 |  | 91.72 | 89.92 |  |  |
| Nucleotide | — |  | — | — |  |  |
| Ligand | — |  | — | 65.72 |  |  |
| **R.m.s. deviations** |  |  |  |  |  |  |
| Bond lengths (Å) | 0.009 |  | 0.008 | 0.007 |  |  |
| Bond angles (°) | 1.153 |  | 1.201 | 1.16 |  |  |
| **Validation** |  |  |  |  |  |  |
| MolProbity score | 2.23 |  | 1.7 | 1.63 |  |  |
| Clashscore | 18.61 |  | 6.9 | 5.61 |  |  |
| Rotamer outliers (%) | 0.15 |  | 0 | 0.44 |  |  |
| **Ramachandran plot** |  |  |  |  |  |  |
| Favored (%) | 92.59 |  | 95.47 | 95.36 |  |  |
| Allowed (%) | 7.41 |  | 4.53 | 4.64 |  |  |
| Disallowed (%) | 0 |  | 0 | 0 |  |  |

### Cryo-electron Microscopy Image Processing

Stack images were exported to cryoSPARC live (v3.3.1 or v4.4.0), where they underwent motion correction and CTF determination. Particles were picked using 2D class averages generated from initial templates, then exported to cryoSPARC [57] for additional processing. Contaminants, broken vaults, and poorly-aligned classes were eliminated following initial 2D classification. Depending on the dataset, between 100,000 and 318,989 particles were used to generate *ab initio* classes that were typically of low quality but useful as templates to sort particles during heterogenous refinement. For the first homogeneous refinement, a low-pass filtered MVP cage map from an earlier dataset served as the initial model, with D39 symmetry imposed. The refined map and previous low-quality *ab initio* volumes were used as inputs for a heterogeneous refinement job to further sort vault cage particles. Particles aligned to the well-resolved MVP cage map were subjected to non-uniform refinement. The resulting map was typically used as the input for an additional round of either homogeneous or non-uniform refinement, in which the program was instructed to optimize the per-particle defocus and per-exposure-group CTF parameters, fit the anisotropic beam magnification, and apply Ewald sphere correction using the correct curvature sign, as determined by running homogeneous reconstruction with both. See Supplementary Figures 2 and 3 for more detail.

### Cryo-electron Microscopy Model Building, Refinement, and Validation

Model building was completed in COOT [58] with starting models including the rat homologue of MVP from a previous crystal structure of the vault cage (PDB accession no. 4V60) and PARP4’s MINT domain predicted by AlphaFold2 [59]. Small molecule coordinates were retrieved from the REFMAC monomer library in COOT. Full length PARP4 was present in the complex, but only a model of the MINT domain could be confidently built into the potential. From the cryo-EM map, it appeared as though there were 78 copies each of MVP, MINT, and bound ligand in the ternary complex. However, based on known space constraints and estimated protein stoichiometries (from SDS-PAGE gels) of approximately 1:2 PARP4:MVP in the recombinant complex, this is likely an artefact of particle averaging and symmetrization.

Therefore, although two regions of the MINT domain had to be deleted in order for the map to accommodate adjacent MVP-MINT models, we maintain that the entire domain should fit into the complex in the absence of artefactual PARP4 symmetry. The final model, containing two copies of MVP related by a 2-fold axis, was refined in real space and validated using PHENIX [60]. See **Table 1** for the details of model statistics.

### Isothermal Titration Calorimetry (ITC)

MVP cage protein was purified mostly as described above, with the exception that the buffer was DTT-free (since it tends to oxidize during injection and distort the ITC baseline).

Sample was concentrated to 320 µl and subjected to a 10 minute spin at 13,000 rpm, at which point the supernatant was transferred to a new tube. The MVP monomer concentration was calculated to be about 50 µM by protein absorbance measurement at 280 nm. A 5 mM solution of NAD*^+^ (*ChemImpex 00229) was prepared in the exact buffer used for MVP SEC. ITC binding experiments were conducted using a MicroCal PEAQ-ITC instrument (Malvern Panalytical).

Following an initial 0.4 µl ligand injection, 18 injections of 2 μl NAD^+^ solution were titrated into buffer (as a control) or MVP at a rate of 2 s μl^−1^ at 150 s time intervals. Stirring speed and experimental temperature were kept at 750 rpm and 25 °C, respectively.

### Gel Filtration Peak Shift Assays

Wild-type and mutant MINT (100 µM) and MVP3-5 (90 µM) constructs were individually injected onto a Superdex 75 increase 10/300 GL column (Cytiva) to establish the peak height and elution volume of each construct. To assess complex formation, wild-type MINT (100 µM) was incubated with wild-type MVP3-5 (90 µM) overnight at 4°C. The mixture was then injected onto the Superdex 75 increase column. This process was repeated with a mixture of the mutant MINT and MVP3-5 constructs, as well as with combinations of each WT and mutant construct. 2x Laemmli loading dye was added to peak fraction samples, which were then boiled, and loaded onto 15% SDS-PAGE gels. Bands were resolved with Coomassie stain.

### Molecular Dynamics Simulations

The all-atom simulation system was built using a dimer of the SPFH domain (residues: 419-646) from the atomic cryo-EM structure of the vault cage with NAD^+^ in the binding pocket. The protein with the NAD molecule was solvated in a water box using the VMD program. Titratable residues of the protein were assigned to their default protonation state at neutral pH. Potassium and chloride ions were added into the water box to make the simulation system electronically neutral and reach a final concentration of 100 mM KCl. The resulting system contained ∼55,600 atoms with orthorhombic box dimensions of ∼70 x 80 x 95 Å^3^. The molecular dynamics simulation was performed using the program NAMD [61]. The CHARMM36 force field with torsional backbone corrections was used for protein, ion, NAD, and the TIP3P model for water. The system temperature and pressure were maintained at 300 K and 1 atm using the Langevin dynamics and the Nose−Hoover Langevin piston method, respectively. The long-range electrostatic interaction was calculated using the particle mesh Ewald (PME) method [62], and the van der Waals interaction was gradually switched off at 10−12 Å. A time step of 2 fs was used. The simulation system was initially minimized for 5,000 steps, and then equilibrated for 50 ns with 3 and 7 restraints being applied to the protein and the NAD molecule, respectively, to restrain the conformation, position and orientation of the protein, and the conformation and relative position/orientation of NAD with respect to the protein. The restraints were applied using the Colvars module, with the collective variables being generated using the Binding Free-Energy Estimator 2 (BFEE2) software [63]. The last 10 ns simulation trajectory (1,000 snapshots) was used to calculate the average pairwise interaction between NAD and each protein residue using NAMD.

### A549 Cell Culture and PARP4 KO Cell Line Generation

A549 cells were cultured in DMEM (MilliporeSigma) supplemented with 10% fetal bovine calf serum (Cytiva), 100 U ml^−1^ penicillin, and 100 µg ml^−1^ streptomycin in a humidified incubator (with 95% air, 5% CO2) at 37 °C. Mutant cells were generated via CRISPR–Cas9 transfection. A Guide RNA targeting exon 1 of PARP4 was annealed into the pSpCas9(BB)-2A- GFP plasmid (PX458, addgene) (Supplementary Table 2). The guide RNA-Cas9 plasmid was transfected into A549 cells using Lipofectamine 3000 (ThermoFisher) and a cell line-specific protocol provided by ThermoFisher (58). Cells were viewed under a fluorescence microscope to verify that transfection had occurred, then trypsinized and pooled for fluorescence assisted cell sorting (FACS) at the University of Chicago Flow Cytometry Core. Each cell that exhibited low-level fluorescence was selected and transferred to an individual well of a 96-well plate. Plates were transferred to a 37°C incubator and kept there overnight, after which the cells were supplemented with fresh media. Single clone cell lines were expanded to larger plates as needed. The genomic DNA of the cell lines that persisted was isolated and genotyped to identify mutants. Western blot assays were performed to verify protein-level depletion (Fig. 5b)

### anti-hMVP sAB selection and purification

Human MVP was expressed and purified as described above with the exception that MVP was purified in amine-free buffer (150 mM NaCl, 50 mM HEPES pH 7.4, 1.5 mM MgCl2, 0.1 mM TCEP-HCl) prior to sAB selection using the phage display library as described previously [64] . MVP was biotinylated and immobilized on streptavidin beads, at which point the beads were washed with selection buffer and subjected to four rounds of phage display panning at increasingly lower concentrations of MVP (200-10 nM). The nineteen sAB candidates that emerged from the final round of clones were subjected to a phage ELISA assay, and the twelve that displayed the greatest affinity and specificity for MVP were subcloned into the RH2.2 expression vector for subsequent purification as documented previously [65]. The sAB with greatest affinity by ELISA (VM1) was selected for coimmunoprecipitation assays.

### Vault Immunoprecipitation and Mass Spectrometry

Three anti-human MVP and control IP biological replicates were conducted per cell line (WT versus PARP4 KO) for a total of twelve samples. WT and PARP4 KO A549 cells were seeded (at 7.5E5 cells per 15-cm plate) three days prior to the experiment. Two 15-cm plates were seeded per condition replicate (for a total of 24 plates). On the day of the experiment, cells were washed with 8 and then 4 ml ice cold PBS, then subjected to lysis with 1.7 ml fresh NETN buffer (150 mM NaCl, 50 mM Tris pH 7.5, 5 mM EDTA, 0.5% NP-40, 1 mM DTT, 1 mM PMSF, 1 complete protease inhibitor tablet (Roche) per 20 ml buffer, and 250 nM ADP-HPD PARG inhibitor). Cells were incubated with NETN buffer at 4°C for 6 minutes, at which point the lysate was collected with a cell scraper and transferred to a 2 ml centrifugation tube. Tubes were vortexed for 30 s and kept on ice for an additional 15 mins. Cell debris and nuclei were pelleted for 8 minutes at 13,000 rpm, after which the lysate supernatant was transferred to a fresh 2 ml tube and placed on ice. Following a Bradford assay to quantify total lysate protein material per replicate, volumes containing 6 mg lysate protein would be used as starting material per coimmunoprecipitation replicate. A total of 55 mg (per replicate) of biotinylated anti-hMVP or anti-GFP (control) sAB were incubated with magnetic streptavidin beads at 4°C. After being washed with NETN buffer three times, the sAB-bead mixtures were added to harvested lysate and incubated in an overhead rotator at 4°C for two hours in order to limit the binding of false positives. Samples were then subjected to a stringent series of eight washes by NETN buffers with varying NaCl concentrations. The wash cycle started with the highest (2X at 500 mM) NaCl concentration and progressed to lower (3X at 300 mM), then lowest (3X at 150 mM) NaCl concentration buffers. All washes were conducted on an overhead rotator at 4°C. Finally, immunoprecipitated proteins were briefly washed with 900 µl ddH2O to eliminate buffer contaminants, then eluted from the streptavidin beads with 82 µl 0.1 M acetic acid, pH 4.5, and incubated at 37°C. Elutions were immediately stored at -80°C for subsequent MS analysis.

Samples were reduced, alkylated, and trypsin-digested overnight. Peptides were eluted and desalted with C18 stagetip, then sent for MS analysis. Data was searched with Proteome Discoverer 3.0 [66], normalized with Perseus 2.0.10.0 [67], and analyzed using Microsoft Excel and R Studio.

## Acknowledgements

We thank Jotham Austin, Tera Lavoie, and James Fuller at the University of Chicago Advanced Electron Microscopy Core, as well as Rui Yan, Xiaowei Zhao, and Zhiheng Yu at the HHMI Janelia Research Campus, for their assistance with cryo-EM data collection. We also thank Virender Kumar, Varun Sharma, and Himanshi Yadav for maintenance of and assistance with University of Chicago Research Computing Cluster resources and the Beagle3 GPU cluster, which is supported by the National Institute of Health (NIH) under the High-End Instrumentation (HEI) grant program award S10OD028655. Isothermal titration calorimetry data were collected with the help of Anchal Sharma, Robel D. Demissie, and Hyun Lee at the University of Chicago-Illinois Biophysics Core. ADP-ribosylation identification mass spectrometry data were collected and analyzed with the help of Lasanthi Jayathilaka at the University of Illinois-Chicago Mass Spectrometry Core, and proteomics data were collected and analyzed with the help of Allen Huff and Samuel Weng at the University of Chicago Proteomics Platform. This work was supported by the NIH under grant number R35GM143052 to M.Z. and by a training grant T32GM144290 to J.E.L.

## Data Availability

Cryo-EM maps of the human vault cage on its own, bound to PARP4, and bound to PARP4 and NAD^+^ have been deposited in the Electron Microscopy Data Bank (EMDB) under accession codes EMD-44953, 44955, and 44957. Atomic coordinates for the above complexes have been deposited in the Protein Data Bank with accession codes 9BW5, 9BW6, and 9BW7, respectively. Additional cryo-EM maps of the human vault cage with NAD^+^, ADP, and the MINT domain of PARP4 have been deposited in the EMDB under accession codes EMD-44954, 44959, and 44960. Other data associated with this manuscript is available upon reasonable request.

## Author contributions

J.E.L. and M.Z. designed the research. J.E.L. performed all laboratory experiments and prepared specimens for cryo-EM data collection. J.E.L. and M.Z. collected and processed cryo-EM data and built and refined atomic models. J.E.L. and M.Z. wrote the manuscript with input from the other authors. R.S. performed MD simulations and processed the results. S.M. and A.A.K. performed and directed synthetic antigen binding reagent selection and validation assays. Y.X. provided plasmids and reagents for protein expression and optimized pull-down protocol for mass spectrometry, and K.R. assisted with protein purification.

