## Supplemental Information for "Structural Insights into the Roles of PARP4 and NAD^+^ in the Human Vault Cage"

Supplementary Information

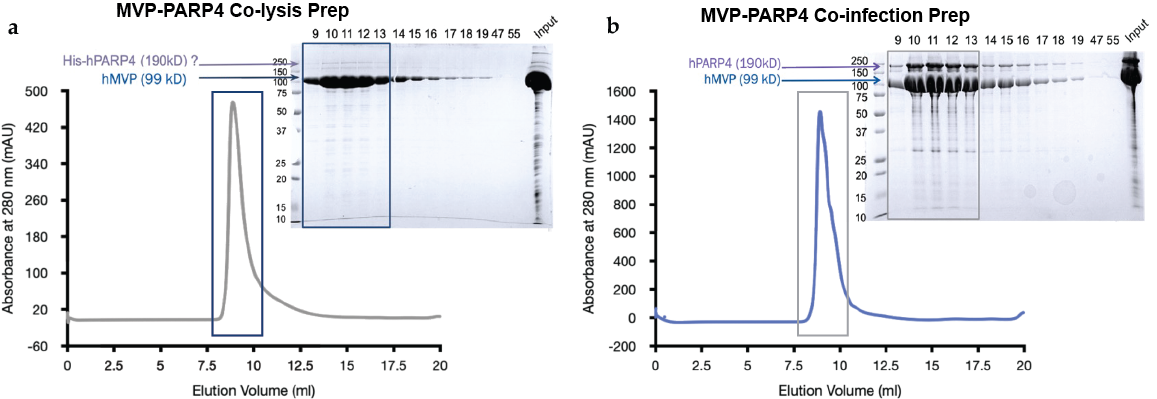

**Supplementary Figure 1. PARP4 is not efficiently incorporated into the vault cage *in vitro* | a**, SEC chromatogram of an attempt to purify the MVP-PARP4 complex via insect cell co-lysis (left) and SDS-PAGE gel (right) corresponding to that peak. **b,** SEC chromatogram of the purified MVP-PARP4 complex following co-infection of insect cells with each vault component’s respective baculovirus (left). The corresponding SDS-PAGE gel (right) indicates that PARP4 can only be efficiently internalized in the MVP cage when both complex components are co-expressed in insect cells.

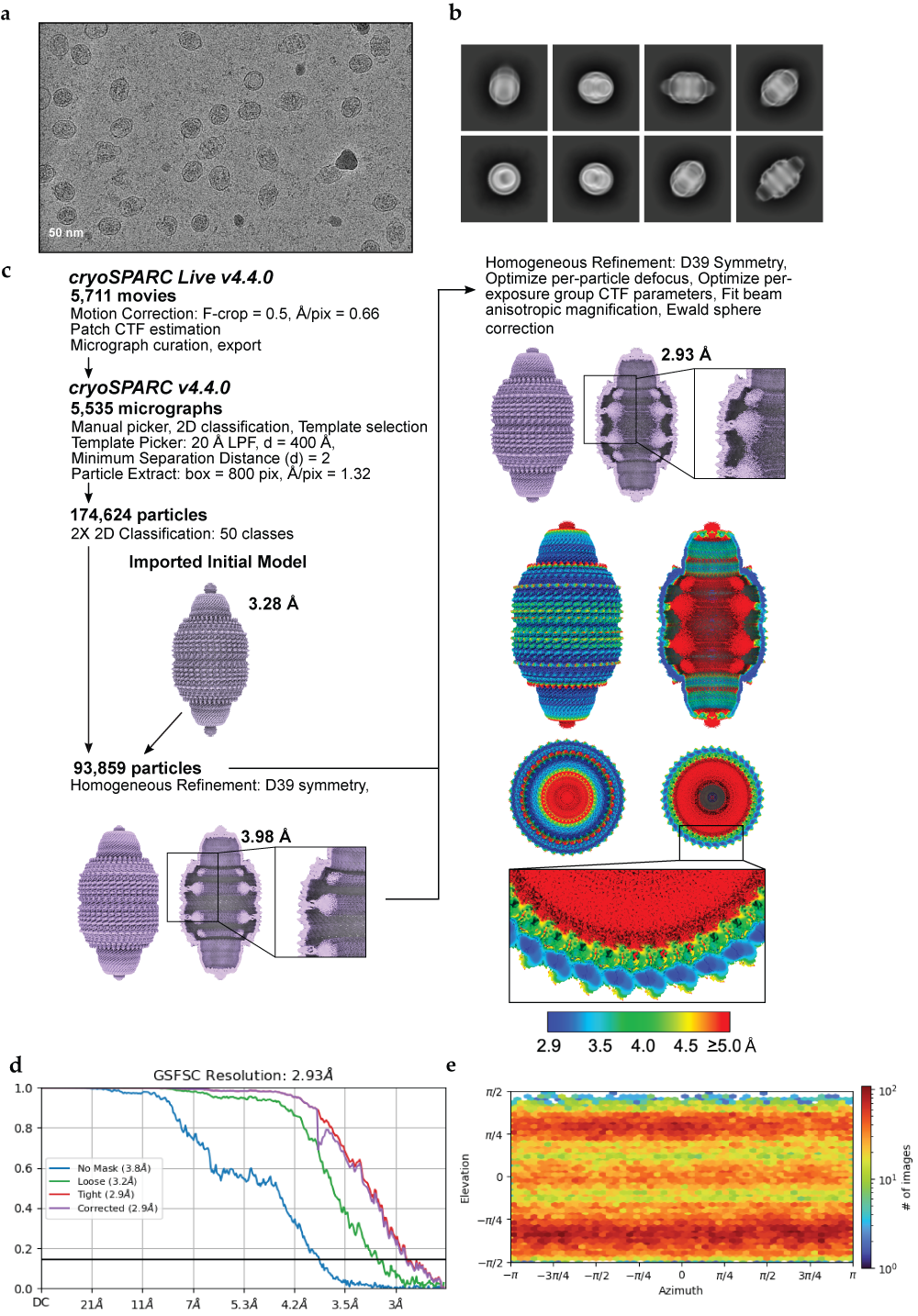

**Supplementary Figure 2. Data processing workflow for the MVP-PARP4-NAD^+^ structure | a,** A representative motion-corrected electron micrograph from the MVP-PARP4-NAD^+^ dataset (n = 5,711), collected on a Titan Krios microscope. **b,** A subset of initial 2D class averages, following template particle picking. **c,** The image processing workflow for MVP-PARP4-NAD^+^. Initial processing was completed using cryoSPARC LIVE v4.4.0. Selected particles were exported to cryoSPARC v4.4.0. Exported particles were subjected to several rounds of homogeneous refinement, beginning with an initial model resolved from a previous dataset. The final map is shown with local resolution values calculated in cryoSPARC v4.4.0 **d,** Fourier Shell Correlation (FSC) curves as calculated in cryoSPARC. **e,** Euler angle distributions.

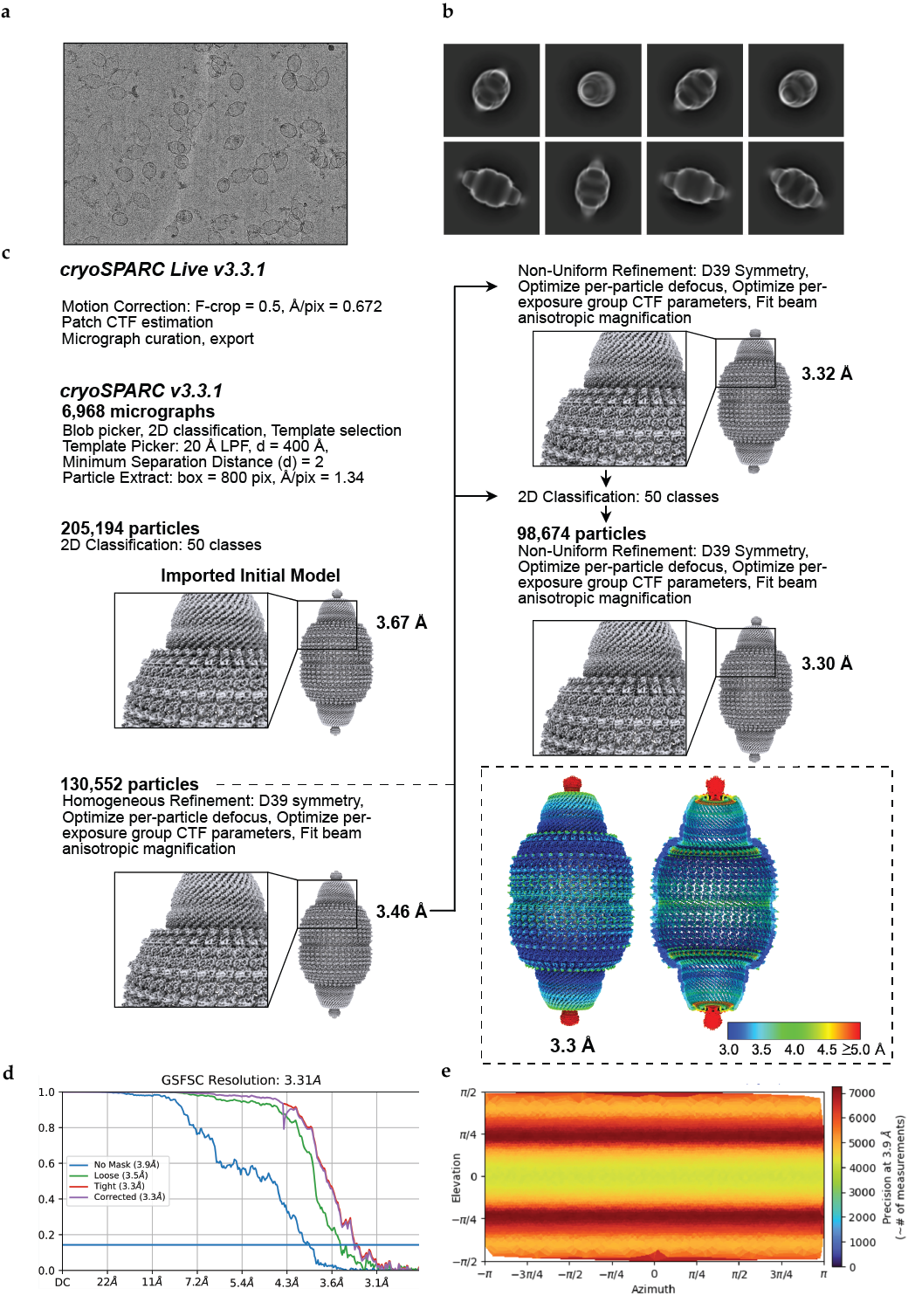

**Supplementary Figure 3. Data processing workflow for the apo-hMVP structure | a,** A representative motion-corrected electron micrograph from the apo-hMVP dataset (n = 6,968), collected on a Titan Krios microscope. **b,** A subset of initial, reference-free 2D class averages, following “blob” particle picking. **c,** The image processing workflow for apo-hMVP. Initial processing was completed using cryoSPARC LIVE v3.3.1. Selected particles were exported to cryoSPARC v3.3.1. Exported particles were subjected to several rounds of homogeneous and non-uniform refinement, beginning with an initial model resolved from a previous dataset. The final map is shown with local resolution values calculated in cryoSPARC v3.3.1. **d,** Fourier Shell Correlation (FSC) curves as calculated in cryoSPARC. **e,** Euler angle distributions.

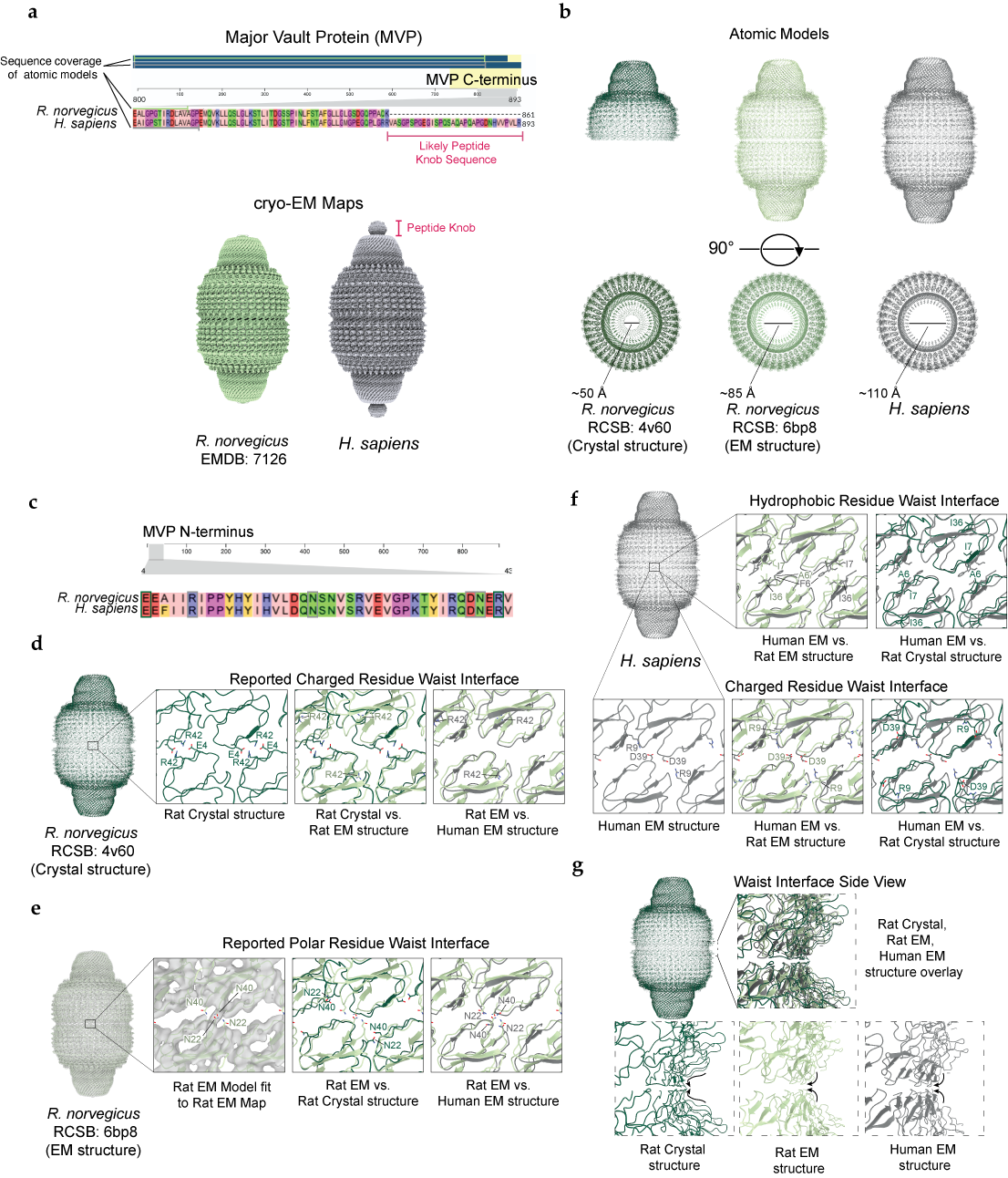

**Supplementary Figure 4. Comparison of our human MVP cage structure with previous rat MVP cage structures | a,** Sequence alignment (upper) of the rat and human MVP C-termini, with residues comprising the human peptide knob underlined in pink. Cryo-EM maps (lower) of the rat MVP cage (light green, left) from a previous study and the human MVP cage (gray, right) from this one. A pink line denotes the peptide knob in the human map**. b,** MVP cage atomic models from: an x-ray crystallography study that solved the structure of half of the rat vault cage (dark green, left), the cryo-EM study that resolved the rat vault cage map shown in a (light green, center), and the human vault cage from the present study (right, gray), viewed from the side (upper) and the top (lower). Top views show openings at the C-terminal caps of the cages. The rat crystal structure model could be built to residue 845, while the rat and human cryo-EM models only extended to residues 815 and 812, respectively. **c,** Sequence alignment of the rat and human MVP N-termini. Residues identified as important for inter-half stability from the original rat vault crystal structure paper are boxed in green and those identified from the present study are boxed in gray, illustrating that all are conserved. **d,** Rat MVP crystal structure, with the full cage (left) generated by retaining the symmetry mate to the top half. Zoomed in images of inter-half interacting residues reported to be key for the inter-half interaction are labeled in dark green (left inset). Overlay with rat EM model (center inset) and overlay between rat and human EM models (right inset) to show the different location of R42 in the EM structures. **e,** Rat MVP model (light green) fit to the corresponding cryo-EM map (gray), which served as the central axis to generate D39 symmetry mates of the monomer model and build the full cage. Magnified view of polar residues that meet at the model’s medial interface, fit to the map (left inset). Overlay with the rat crystal structure model (center inset) and human EM model (right inset) to highlight greater similarity between EM structures. **f,** Human MVP structural model (gray). A dimer model of medially symmetric MVP monomers was fit to the cryo-EM map, which served as the central axis to generate C39 symmetry mates and build the full cage. The previously discussed residues driving inter-half interactions are labeled in the insets, with the rat crystal and EM structural models overlaid to highlight differences. The rat EM structure is broadly similar to the human but with an alanine in place of F6 (left, upper panel) and an R9 rotamer positioned further from the waist interface, such that it clashes with adjacent MVP monomers (center, lower panel). The rat EM and crystal structures once again differ significantly (right panels). **g**, Overlay (inset, side view) of all three models, highlighting the significant differences at the waist in the crystal (lower left) versus EM (lower center and right) structures.

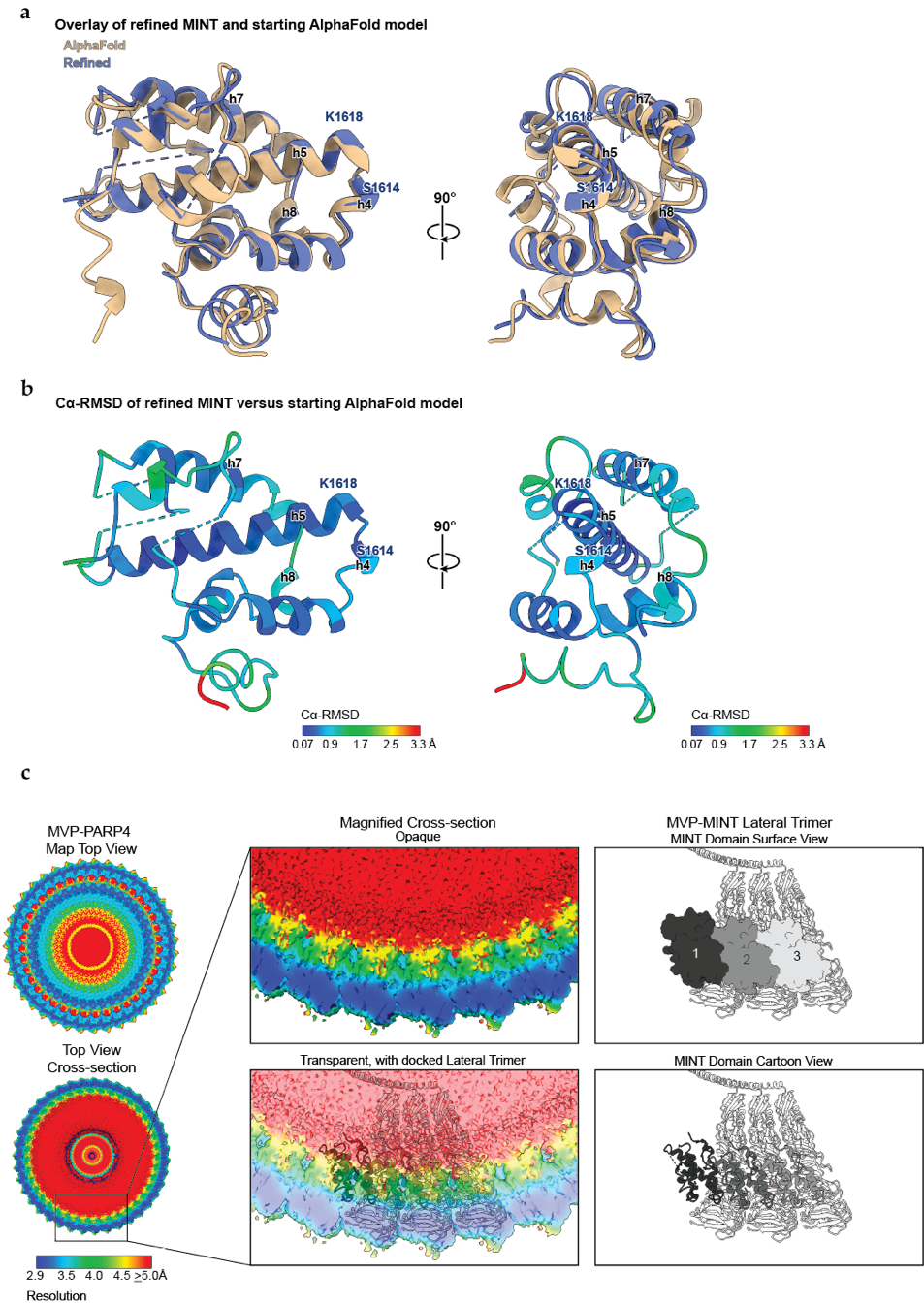

**Supplementary Figure 5. MINT Domain Model Building | a,** Overlay of the starting Alphafold2 model of the MINT domain (tan) and the final refined version (blue). Residues that were deleted from the final MINT model are represented by dashed lines. Helices and residues critical for MVP interaction are labeled. **b,** Per-residue alpha-carbon RMSD (Å) of the aligned Alphafold2 and refined MINT models. Helices and residues critical for MVP interaction are labeled. **c,** Top view (upper left) and cross-sectional top view (lower left) of the MVP-PARP4 cryo-EM map, colored by local resolution. Magnified view of the map cross-section (upper middle) and transparent view of the same segment of the map, with three, lateral MVP-PARP4 atomic models docked into it (lower middle). View of the models at the equivalent position with the MINT domain shown in surface view (upper right) and cartoon view (lower right) to highlight differences in register between the MINT domains and MVP chains. Each MINT domain is colored in a successively lighter gray to distinguish between them.

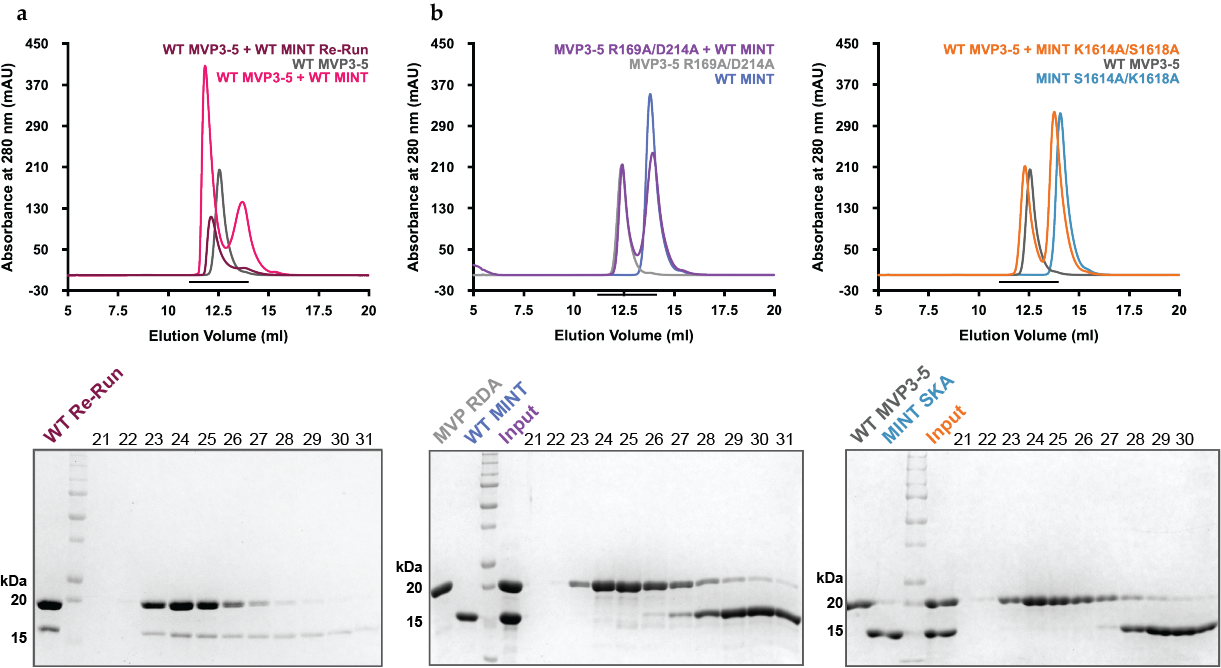

**Supplementary Figure 6. Supplementary SEC complex traces | a,** SDS-PAGE gel (upper) showing the comigration of the previously purified WT MVP3-5 and MINT complex, re-run over a Superdex 75 column. Overlay of traces corresponding to WT MVP3-5 (gray), the original WT MVP3-5 + WT MINT complex run (pink), and the re-run of the complex fractions (dark red). **b,** SDS-PAGE gels (upper) showing lack of co-migration between each MVP3-5 and MINT double mutant and the WT construct of the other species, following their co-injection over a Superdex 75 column. Corresponding size exclusion chromatograms beneath each gel show individual traces of MVP3-5 (gray) and MINT constructs (blue), measuring their absorbance at 280 nm. Individual traces are overlaid with complex traces following co-incubation of mutant MVP3-5 with WT MINT (purple, left) and WT MVP3-5 with mutant MINT (orange, right). Black bars denote the area over which fraction samples were collected.

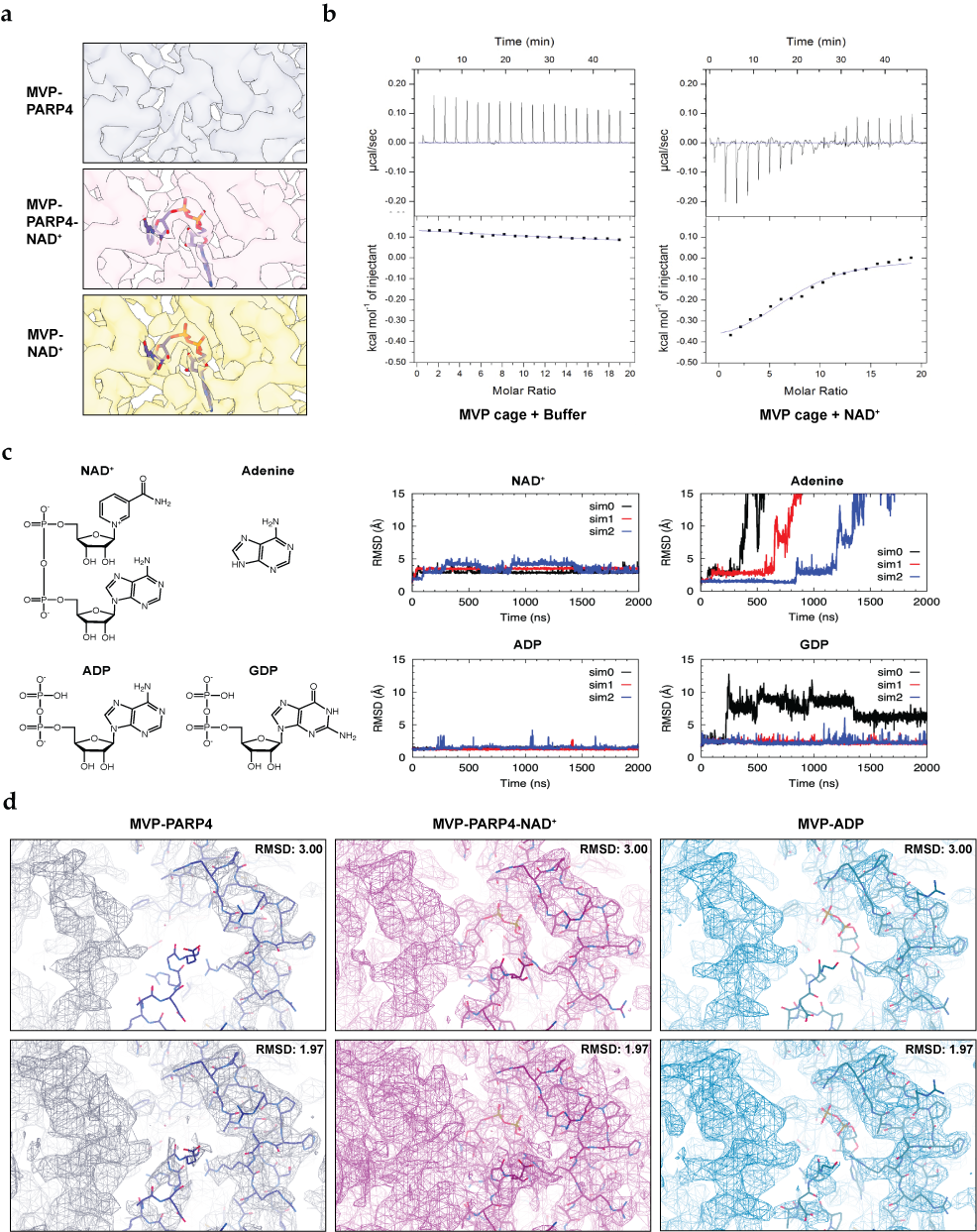

**Supplementary Figure 7. Interactions between MVP and Nucleotides | a,** Cryo-EM maps of the MVP-PARP4 (upper), MVP-PARP4-NAD^+^ (middle), and MVP-NAD^+^ (lower) complexes, focused on MVP’s ligand binding site. NAD^+^ is docked into the additional potential found in the ligand-bound maps. **b,** Isotherms of buffer (center) and NAD^+^ (right) injected into a solution of the MVP cage. 2 µl titrations of 5 mM NAD^+^ were injected 18 times (following an initial 0.4 µl injection) into a cell containing MVP purification buffer, inducing a dose-independent endothermic response (center). Identical concentrations and volumes of NAD^+^ were injected into a solution of the MVP cage complex (with MVP monomer concentration of 50 µM), inducing a dose-dependent exothermic response (right). Lines were fit to normalized ITC data (below). First injection values were disregarded, and the MVP + NAD^+^ binding isotherm was reference-subtracted, with adjacent averaging smoothing applied. **c,** Chemical structures (left) of four ligands simulated in the NAD^+^-binding pocket. A 2 μs RMSD simulation of NAD^+^ in our cryo-EM-derived model of the MVP binding pocket (upper left panel) was performed in triplicate, with each trial trace shown in a different color. Small RMSD fluctuations (<5 Å) indicate base-flipping, while larger deviations suggest that the metabolite has left the binding pocket. Identical simulations (remaining three panels) of adenine, ADP, and GDP suggest that a purine base with ribose and diphosphate moieties is sufficient for stable occupancy of a nucleotide in MVP’s NAD^+^-binding pocket. **d,** Cryo-EM maps (mesh) of the MVP-PARP4 (left), MVP-PARP4-NAD^+^ (middle), and MVP-ADP (right) complexes, focused on MVP’s ligand binding site, and visualized in Coot at equivalent RMSD values: 3.00 (upper) and 1.97 (lower).

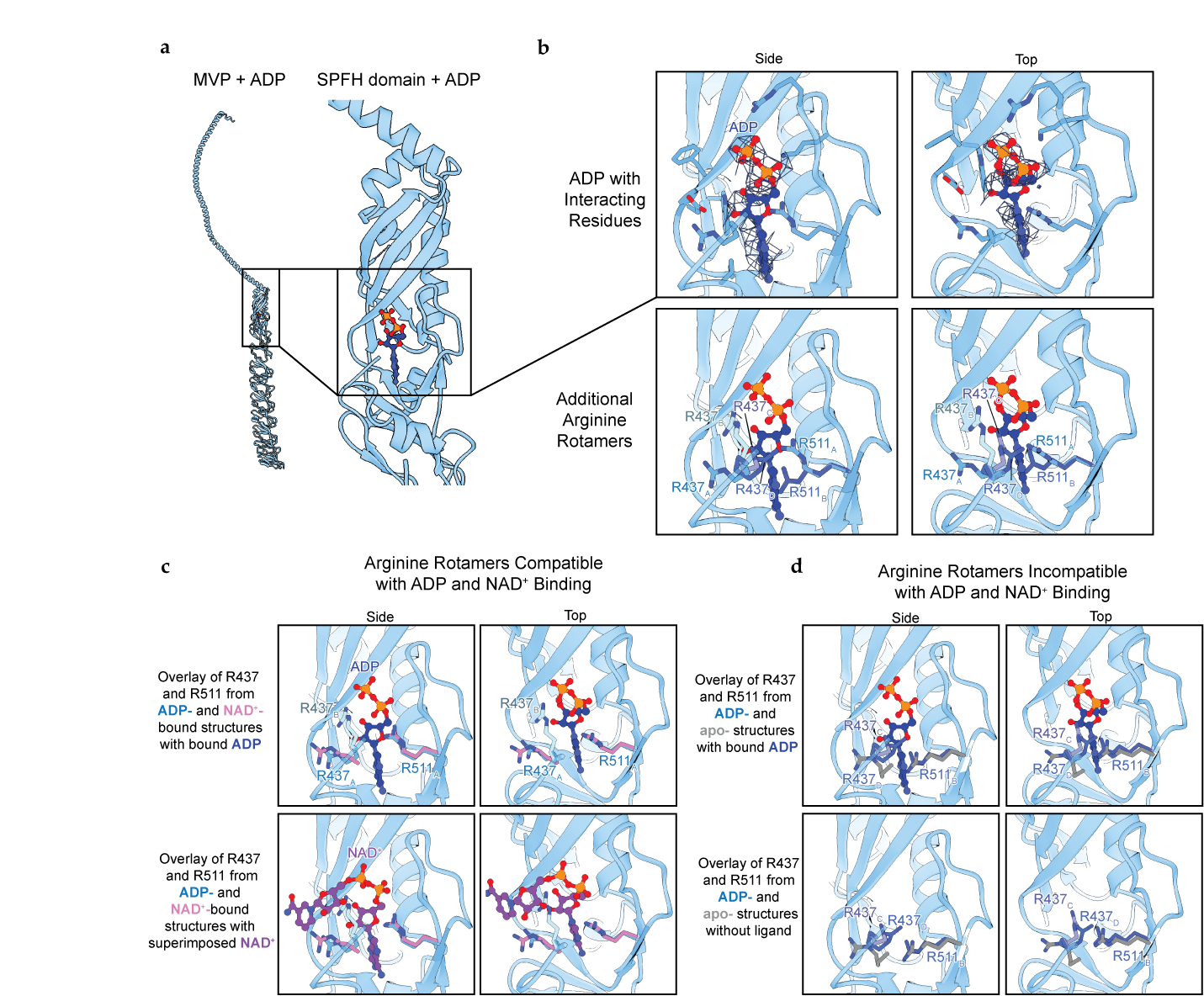

**Supplementary Figure 8. ADP Occupancy of the MVP Ligand Binding Site | a,** Atomic model of the MVP monomer in the presence of ADP (left) and magnified view of ADP in the SPFH domain’s ligand binding site (inset, right). **b,** Side (left) and top (right) views of MVP’s ligand binding site with zoned cryo-EM potential around bound ADP (dark blue) shown as mesh. Side chains of previously identified ligand binding resides shown in stick view (upper). Side (left) and top (right) views of the ligand binding site with an overlay of all rotamer conformations of R437 and R511 that are supported by the cryo-EM potential. Rotamers R437A-B and R511A can accommodate bound ADP, while R437C-D and R511B cannot. **c,** Side and top views of overlaid R537 and R511 rotamers from the MVP-ADP (blue, cyan) and MVP-NAD^+^ (pink) structures that support ligand binding. Bound ADP (upper) and bound NAD^+^ (lower) are both shown occupying the pocket to illustrate the ability of the R437B rotamer to accommodate ADP but not NAD^+^. **d,** Side and top views of overlaid R537 and R511 rotamers from the MVP-ADP and apo-MVP structures that are incompatible with ligand binding. The pocket is shown with bound ADP (upper) and in its empty state (lower) to illustrate the steric hindrance imposed by R437C-D and R511B.

| ADP-ribosylated Hits (from Ref. 49 Ayyappan et al.) | Hits that Bind ADP-ribose (from Ref. 47, Gagné et al. 2012 and Ref. 48, Wright et al. 2016) | Hits that Bind NAD (from Ref. 43, Duarte-Pereira, et al.) |
| --- | --- | --- |
| AKR1B1 | AKR1B2 | AKR1B1 |
| AKR1C2 | ANXA1 | AKR1C2 |
| AKR1C3 | ANXA2 | AKR1C3 |
| ALDH1A1 | DAZAP1 | ALDH1A1 |
| ANXA1 | DDX56 | PARP4 |
| ANXA2 | EZR | SLC25A1 |
| CASP14 | HIST1H3A | TPI1 |
| CEP295 | HSP90AA1 |  |
| CFAP20 | HSP90AB1 |  |
| DARS2 | IPO7 |  |
| DAZAP1 | KPNB1 |  |
| DDX56 | NAP1L1 |  |
| EIF1B | MRPL15 |  |
| EZR | MVP |  |
| FIP1L1 | PIP |  |
| GMPPA | PPIA |  |
| HNRNPDL | PRDX1 |  |
| HSP90AA1 | PUM1 |  |
| HSP90AB1 | RACK1 |  |
| IPO7 | RUVBL2 |  |
| KPNB1 | SLC25A1 |  |
| KRT78 | SMC6 |  |
| LRRFIP2 | TAGLN2 |  |
| MICAL2 | TPI1 |  |
| MAP7 | TXN |  |
| MRPL15 | UBAP2L |  |
| MVP |  |  |
| NAP1L1 |  |  |
| NBR1 |  |  |
| NME4 |  |  |
| NOP53 |  |  |
| NSMCE1 |  |  |
| NSMCE3 |  |  |
| PARP4 |  |  |
| PIP |  |  |
| PPIA |  |  |
| PRDX1 |  |  |
| PUM1 |  |  |
| PUM2 |  |  |
| RACK1 |  |  |
| RUVBL2 |  |  |
| S100A10 |  |  |
| SLC25A1 |  |  |
| SMC5 |  |  |
| SMC6 |  |  |
| TAGLN2 |  |  |
| TEP1 |  |  |
| TPI1 |  |  |
| TXN |  |  |
| UBAP2L |  |  |
| VCP |  |  |
| WDR11 |  |  |

**Supplementary Table 1. MS Hits that are ADP-ribosylated, known to bind poly(ADP-ribose), and known to bind NAD^+^.** Cells are filled in with blue, purple, or pink, corresponding to proteins in the WT, shared, and KO hits datasets, respectively.

**Supplementary Table 2. Primer sequences for constructs used in the research**

| Construct | Forward/S Primer (5'-3') | Reverse/AS Primer (5'-3') |
| --- | --- | --- |
| pVL1393-hMVP | CCACCATCGGGCGCGATGGCAACT  GAAGAGTTCATC | CTAGAAGGTACCCGGTTAGCGCAG  TACAGGCACC |
| pVL1393-hPARP4 | CCGTCCCACCATCGGGCGCGATGG  TGATGGGAATCTTTGCAAATTG | CTAGAAGGTACCCGGTTAGCCTTGA  CTGTAATGGAGGACTC |
| pET47b-MVP3-5 | GGACCCGGGTACCAGGGGGAGGTG  CTGGAAAAGG | GGCCTGTACAGAATTCGTTAGGTGA  TGGGCACAACCCC |
| pET47b-MINT | TCAGGGACCCGGGTACCAGGTGTG  CATACAACACTGGC | GGCCTGTACAGAATTCGTCAGCCTT  GACTGTAATGGAGGAC |
| MVP3-5 R169A | TCATCGCGCAGAACCAGGCTC | TTCTGCGCGATGATGGTGGCC |
| MVP3-5 D214A | TTCTGGCTTTGGTGGACGCC | CCAAAGCCAGAACCTCCTCAAACAC |
| MINT S1614A/K1618A | GTGTTCAAAGCACTGATGGCAATGG  ATGAC | GCCATCAGTGCTTTGAACACTATTC  CCTC |
| MINT K1618A | CACTGATGGCAATGGATGACGCTTCT | CATCCATTGCCATCAGTGATTTGAA  CACTA |
| PARP4 KO sgRNA | CACCGCTGGGTTTGCAATATGAACG | AAACCGTTCATATTGCAAACCCAGC |
